# Microlenses fabricated by two-photon laser polymerization for cell imaging with non-linear excitation microscopy

**DOI:** 10.1101/2022.11.17.516871

**Authors:** M. Marini, A. Nardini, R. Martínez Vázquez, C. Conci, M. Bouzin, M. Collini, R. Osellame, G. Cerullo, B.S. Kariman, M. Farsari, E. Kabouraki, M.T. Raimondi, G. Chirico

**Affiliations:** Department of Physics, Università degli Studi di Milano-Bicocca, Piazza della Scienza 3, 20126, Milan, Italy; Institute for Photonics and Nanotechnologies (IFN), CNR, Piazza L. da Vinci 32, 20133 Milan, Italy; Department of Chemistry, Materials and Chemical Engineering “Giulio Natta”, Politecnico di Milano, Piazza L. da Vinci 32, 20133 Milan, Italy; Department of Physics, Politecnico di Milano, Piazza L. da Vinci 32, 20133 Milan, Italy; FORTH/IESL, N. Plastira 100, 70013, Heraklion, Greece

**Keywords:** 3D micro scaffolds, two-photon polymerization, microlenses, two-photon imaging, confocal microscopy, SZ2080

## Abstract

Non-linear excitation microscopy offers several advantages for in-vivo imaging compared to conventional confocal techniques. However, tissue penetration can still be an issue due to scattering and spherical aberrations induced on focused beams by the tissue. The use of low numerical aperture objectives to pass through the outer layers of the skin, together with high dioptric power microlenses implanted in-vivo close to the observation volume, can be beneficial to the reduction of optical aberrations. Here, we develop and test on fibroblast cell culture plano-convex microlenses to be used for non-linear imaging of biological tissue. The microlenses can be used as single lenses or multiplexed in an array. A thorough test of the lenses wavefront is reported together with the modulation transfer function and wavefront profile. We could retrieve magnified fluorescence images through the microlenses coupled to commercial confocal and two-photon excitation scanning microscopes. The signal-to-noise ratio of the images is not substantially affected by the use of the microlenses and the magnification can be adjusted by changing the relative position of the microlens array to the microscope objective and the immersion medium. These results are opening the way to the application of implanted micro-optics for optical in-vivo inspection of biological processes.

## 1. Motivation/introduction

*In vivo* observation of tissue is desirable in many fields, from immunology^[1,2]^ to the assessment of the foreign body reaction as a test of immunogenicity to biomaterials.^[3,4]^ To date, different kinds of imaging windows have been developed and used^[5]^ in combination with fluorescence microscopy: the cranial imaging window, the dorsal skinfold chamber, the mammary imaging window, and the abdominal imaging window.^[6]^ The dorsal skinfold chamber is one of the most widely used devices for *in vivo*^[7]^ observation of the reaction to biomaterials implanted sub cute (as required by the ISO 10993-6 norm).^[8]^ This chamber typically comprises two titanium frames fixing the extended dorsal skin in the back’s midline, it is typically 4 × 3 cm wide and weighs a few grams.^[7]^ Weight and skin stretching may lead to restricted breathing, immobilization, and pain. The severity of any dorsal skinfold chamber experiment is considered at least moderate, and its use for extended observations (more than 10 days) has been questioned in the view of the 3R’s (refinement, reduction, replacement) principles.^[9]^ Apart from the standard bulky titanium chambers, more advanced plastic chambers made from polyetheretherketone (PEEK) have been recently used,^[10]^ which sensibly reduces the animal distress. Cranial and abdominal imaging windows have lower impact on the animals. However, there is still the need to develop methods to reduce the impact of the implant of these types of windows on the laboratory animal health and to limit the changes induced in the animal behavior, therefore ensuring the medical relevance of the observations.

We recently proposed a miniaturized (500 μm × 500 μm × 100 μm) imaging window, named Microatlas, which incorporates an array of micro-scaffolds, able to guide the regeneration of tissue at the interface with the implanted material.^[11]^ Although the Microatlas has been validated up to now in embryonated chicken eggs,^[11]^ it can also be implanted sub cute in mice, in which case non-linear excitation microscopy would be the best choice for direct imaging. Non-linear excitation microscopy with near infrared (NIR) laser sources can mitigate the penetration issues in tissues^[12–14]^ by reducing the effect of scattering on the Point Spread Function. However, the reduction of scattering with wavelength is decreasing with a lower than fourth power law in animal tissues,^[15]^ and the spatial resolution of images in deep tissue is also limited by spherical aberrations. These beam aberrations arise simply from the propagation of the high numerical aperture beam through the different layers of the animal skin. Instead of imaging implanted scaffolds by means of the long working distance and high numerical aperture objective of an external microscope, our long-term purpose is to develop a set of microlenses that will be coupled directly to the micro scaffolds, implanted *in vivo*, and used to enable non-linear imaging of the tissue regenerated within the micro scaffolds, by focusing collimated beams right at the site of observation. Structuring of these microlenses directly on top of the micro scaffolds would then reduce the tissue-induced spherical aberrations and allow longitudinal studies of tissue regeneration *in vivo*. In order to image micro scaffolds extending *≃* 100 μm along the optical axis and some 300-500 μm, in the focal plane, we would need to fabricate microlenses with high curvature, large size (semi-diameter at the basis) and smooth surface, and this is a particularly demanding task. This is exactly our goal here, coupled to the need to satisfy the main requirement of non-linear excitation microscopy. For non-linear excitation, the optical design requirements are mainly high numerical aperture (NA) and low group velocity dispersion (GVD), while the main biophysical requirement is to overcome the spherical aberrations due to the tissue inhomogeneity and the tissue scattering. The NA affects the fluorescence signal in at least two ways: by affecting the average intensity at the focal position and by determining the amount of out of focus background. The dependence in both cases is highly nonlinear, ≃ *NA*^4^ and ≃ *NA*^*2*^,^[14,16]^ respectively, making the requirement on the NA stringent for imaging. GVD affects the pulse duration *τ* and, as a consequence, the in-focus signal through the duty cycle, *d*_*c*_ = *τf*_*R*_. The pulse broadening depends on the type of material and on the length of the propagation in the material. Microlenses are relatively thin and should not greatly affect the pulse width that depends on the GVD and the propagation length in the material. Spherical aberrations induced by the light propagation through the tissue can be limited by increasing the size of the microlens with respect to the focal length. These considerations indicate high numerical aperture and large size of the lenses as the most relevant parameters for the design.

Moreover, the fabricated microlenses should have low auto-fluorescence^[17]^ in order not to reduce the S/N of the images, particularly under multi-photon excitation. Among additive manufacturing (AM) techniques, the only suitable technology to achieve this goal is two-photon polymerization (2PP).^[18–22]^ Once the optical design is defined and tested, the solution for a widespread use of the implantable micro-optics would be the use of high throughput techniques like nano-imprint lithography (NIL), starting from high resolution molds fabricated by 2PP.

Microlenses have already been developed and fabricated by 2PP and tested for transmission imaging. They differ mainly for the numerical aperture of the microlens, the size and the 2PP fabrication protocol. From the very first study in this field,^[23]^ attention was drawn to the optimization of the optical quality of the surface, by reducing the fabrication thickness,^[24]^ or by changing the fabrication scanning path.^[23]^ In this work,^[23]^ the microlenses, also in the form of an array, had a numerical aperture 0.03 < *NA* < 0.12 and were tested for conventional transmission imaging only. The fabrication algorithm was based on circular scanning with deterministic access of the starting angle. More recently, the optical characterization has encompassed the study of the Modulation Transfer Function (MTF), which is an indirect measure of the aberrations. For example, the group of Giessen^[25]^ fabricated multi-microlens systems, and characterized them thoroughly for axial and non-axial aberrations. They corrected the spherical aberrations by employing tall multi-elements lenses (about 130 *μm* tall and about 120 *μm* in size) with an effective numerical aperture of about 0.3, one of the largest ever fabricated. This approach, however, could not be easily translated to high throughput fabrication, for example by means of NIL. In a complementary study, the same group achieved the fabrication of free-form optical components directly on top of optical fibers for endoscopy.^[26]^ In the same field of endoscopy, Li et al.^[24]^ demonstrated the feasibility of 2PP of microlenses for optical coherence tomography (OCT) probes. The numerical aperture was not stated, but the optical resolution ≃ 20 *μm* indicates a low value of NA.

To our knowledge, no attempt has been made to fabricate microlenses that can be used under the demanding signal/noise ratio conditions of fluorescence imaging or in the even more demanding case of non-linear excitation microscopy of biological samples. These experimental conditions are particularly relevant for the development of in-vivo biomaterial test protocols that could replace the existing ISO-10993-6 norm,^[8]^ based on ex-vivo histological analyses, with label-free in-vivo microscopy. This goal can be achieved by means of “high-IR” (i.e. excitation wavelength in the range 1300 nm to 1600 nm with pulse energy of about 10 nJ) non-linear imaging, as recently proposed.^[27]^

The present work is therefore specifically devoted to the design, the fabrication and validation of microlenses for two-photon excitation microscopy, reaching large values of the NA over diameters of the lenses of a few hundreds of micrometers. For their fabrication, we choose 2PP, a technology that allows us to reproduce arbitrary surface shapes, with optical quality, thanks to its intrinsically high resolution. In order to fabricate such relatively high volume microlenses, and endeavoring to reduce fabrication time, we found it essential to use a micrometer 2PP voxel for the fabrication of the lens outer shell and a combination of laser two-photon and UV polymerization of the photoresist. We describe methods and protocols for their fabrication and for their experimental testing, both in terms of the optical response functions, and in terms of their direct application on two-photon fluorescence cell imaging. We demonstrate that these microlenses can be used to obtain non-linear excitation fluorescence images on cells, opening the way to their use as implantable devices for the direct inspection of cellular dynamics *in vivo*^[11]^ or in integrated microfluidic setups for the *in vitro* monitoring of complex biosystems, such as organoids, directly in the organoid bioreactor.

## 2. Results and discussion

### 2.1 Microlenses design

The microlenses imply the use of collimated (or almost collimated) light. This is motivated by the rationale of this study, i.e. the use of a microlens or an array of microlenses right at the site of observation, typically an implanted micro scaffold, collecting and focusing a collimated beam.

The skin of a laboratory animal, like a rodent, is composed of several layers,^[28]^ partly made of muscle and partly of adipose tissue. Sebaceous glands and hair follicles, being obstacles larger than the light wavelength, give rise to complex diffraction patterns and also to beam attenuation. Instead, smaller components of the tissue give rise to diffuse scattering, whose effect on imaging can be reduced by adopting a recently developed heterodyne measurement and phase-conjugation correction of the scattered field.^[29,30]^ Apart from these correction strategies, which are not the subject of this work, we notice that different skin layers are endowed with different refractive index values. In fact, the refractive index slightly changes among various tissues, being approximately equal to^[15]^ *n* ≃1.38-1.39 at *λ* ≃ 630 *nm* with the exception of adipose tissue that has *n* ≃ 1.42. The refractive index of the tissues decreases substantially with increasing wavelength^[15]^ (**Fig.SI1**) according to a power law ≈ *λ*^−1.5^ over the range 400 *nm* < *λ*< 700 *nm*, passing from *n* ≃ 1.49 at *λ* = 400 *nm* to *n* ≃ 1.40 at *λ*= 700 *nm*. No data are available yet in the near infrared region typically used for non-linear excitation (*λ*≥ 800 *nm*). However, from the data in ref.[15] we can estimate for the muscle *n* ≃ 1.392 at *λ* = 800 *nm* (**Fig.SI1**). As a matter of fact, microlenses implanted subcute in mice will be probably embedded in the “panniculus carnosus”, similar to muscle, denser but with a lower refractive index than fat. All these considerations imply that the microlenses will be working in an environment having a refractive index about 30% higher than that of air, where lenses are typically tested in optical studies. Therefore, larger curvature of the optical surfaces or even a larger refraction index should be used for the fabrication of our microlenses.

An example of the spherical aberrations observed with high numerical aperture optics in an inhomogeneous medium is simulated in **Fig. 1A**. The propagation medium is either pure water (refraction index *n*_1_ ≃ 1.34 at *λ* = 0.8 *μm*) or composed of two slabs of materials of thickness 8 *mm* < *d*_1_ < 10*mm* (*n*_1_ ≃ 1.34) and *d*_2_ = *f*_FFL_ − *d*_1_ (with refractive index *n*_2_ = 1.45; *f*_*FFL*_ is the front focal length). The test was performed on a plano-aspherical lens, 25.4 mm in size with curvature radius 12.6 mm (see **SI1** for details), optimized for propagation in water and a pupil size of 10 mm. The aberrations induced on the wavefront when increasing the pupil size and/or changing the composition of the propagation medium are primarily spherical and can be characterized by the Zernike polynomial^[31]^ *Z*_4_^0^. The *Z*_4_^0^ amplitude, *a*_4,0_, depends steeply on the pupil size for size larger than 10-15 mm (**Fig.1B**). The spherical aberration coefficient increases more steeply for the propagation in the inhomogeneous medium (*d*_1_ = 8 *mm*) than in pure water (**Fig.1B**) and arises from the mismatch in the refractive indices in the propagation medium as indicated by the linear dependence of *a*_4,0_ on the refractive index *n*_2_ (**Fig.1A**) and on the thickness *d*_2_ of the high refractive index slab (**Fig.1B**, inset). The possibility to obtain the tight focusing of a Gaussian laser beam needed for non-linear excitation microscopy is therefore limited by the propagation in inhomogeneous media. This drawback can be overcome by the use of microlenses implanted close to the observation field.

**Figure 1.**
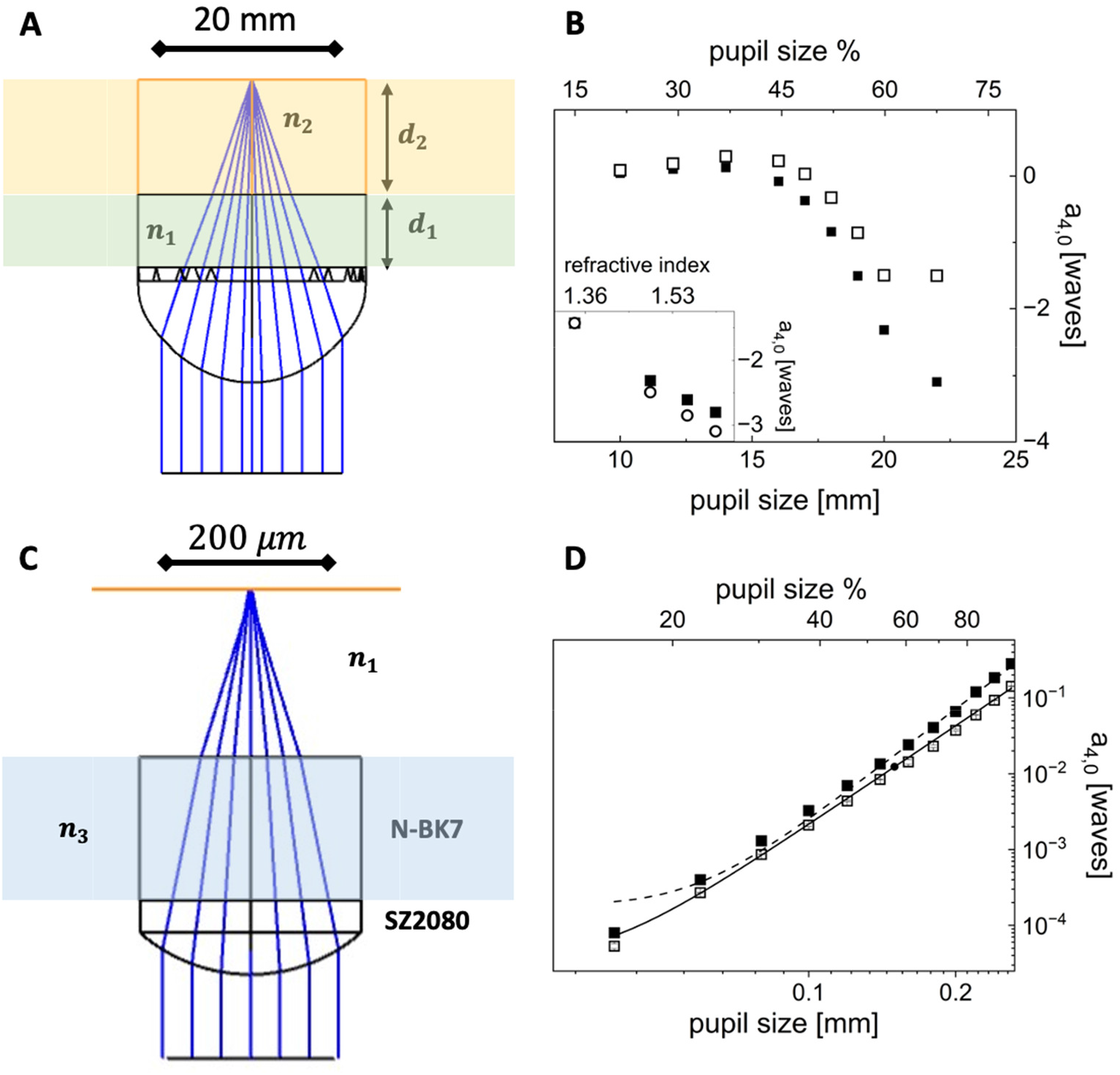
Simulation of the spherical aberrations arising on highly collimated beams when passing through an inhomogeneous sample (*λ*= 0.8 *μm*). **A:** sketch of a macroscopic plano-aspherical lens working in an inhomogeneous medium (refractive indices are *n*_1_ = 1.34 and *n*_2_ = 1.45, 8*mm* < *d*_1_ *<* 10 *mm, d*_2_ = *f*_*FFL*_ − *d*_1_). **B**: spherical aberration coefficient *a*_4,0_ for the lens in panel **A** as a function of the pupil size for propagation in homogeneous (open squares) or inhomogeneous medium (filled squares, *d*_1_ = 8 *mm*). Inset: dependence of *a*_4,0_ as a function of the *n*_2_ refractive index (*d*_1_ = 8*mm* open squares; *d*_1_ = 10*mm* filled squares). **C**: diagram of a single plano-convex microlens (diameter = 264 *μm*) fabricated on a N-BK7 glass substrate (170 *μm* < thickness < 230 *μm*). Tissue is approximated with sea water (*n*_1_ = 1.34). **D**: spherical coefficient *a*_4,0_ for the lens in panel **C** as a function of the microlens pupil size, *p*, in air (filled squares) and in water (open squares). The solid and the dashed lines are the best fit to the data to a power law *a*_4,0_ ∝ *p*^−4.3±0.3^.

Aiming at the development of methods and protocols for rapid prototyping of microlenses with the optical quality needed to prime non-linear excitation, we focus here on the simplest case of a refractive lens, i.e. a plano-convex lens with a pedestal, fabricated on top of a supporting borosilicate glass slide (thickness ≃ 170 *μm*, **Table I**). The plano-convex lens is a good approximation of the shape that theoretically minimizes the spherical aberrations for collimated light.^[32]^ The geometrical design parameters of the lens are reported in **Table I** together with a sketch of the microlens and the location of its principal planes.

**Table I.** geometrical parameters of the plano-convex microlens.

| surface | curvature radius [$\mu\text{m}$] | thickness [$\mu\text{m}$] | material | refractive index [$\lambda = 0.8\text{ }\mu\text{m}$] | radius [$\mu\text{m}$] | sketch |
| --- | --- | --- | --- | --- | --- | --- |
| 1 | 200.0 | 50.0 | SZ2080 | $n = 1.46^*$ | 132.0 | |
| 2 | ∞ | 38.0 | SZ2080 | $n = 1.46^*$ | 132.0 | |
| 3 | ∞ | 170.0 | N-BK7 | $n = 1.52$ | 132.0 | |

Geometrical parameters for a plano-convex lens micro-fabricated in SZ2080. The effective front focal length in air is $f_{FFL} \simeq 164\text{ }\mu\text{m}$ . The two principal planes ( $\Pi$ , dashed lines in the sketch) lie at the vertex V1 and $35\text{ }\mu\text{m}$ below surface 2. \*from ref.<sup>[33]</sup>.

An example of the design of a plano-convex microlens with diameter 264 μm is reported in **Fig.1C** for axial illumination. This microlens allows wide-field imaging with a field of view of about 250 μm (see **SI5, “Field of view of microlenses”**). In order to image wider fields of view, these microlenses could be assembled in a microarray.

The beam focused by the microlens is slightly aberrated even by using the maximum pupil size ≃ 260 *μm*, as can be judged from the intensity profile of a Gaussian beam simulated at the microlens focal plane (**Fig.SI2.2B**) where we can estimate a beam waist (1/e^2^ radius) of (1.20 ± 0.03) *μm* at *λ* = 0.8 *μm* (**Fig.SI2.2**), a value only 2-3 times larger than the one typically used for two-photon excitation in biological samples. The aberrations on the wavefront can be quantified at the focal plane of the microlens (the plane of the circle of minimum confusion) in terms of the Zernike polynomials: we find again that the major contribution for axial propagation is the spherical one corresponding to the Zernike polynomial^[31]^ *Z*_4_^0^. The a_4,0_ coefficient increases steeply as a power law with exponent ≃ 4 (**Fig.1D**), as expected for the thin lens treatment.^[32]^ The spherical aberrations resulted slightly larger when the microlens is focusing in air (**Fig.1D**, filled symbols) rather than in water (**Fig.1D**, open symbols), due to index matching at the exit pupil.

### 2.2 Fabrication and characterization by scanning electron microscopy (SEM)

Following the simulations reported in the previous section, we decided to fabricate plano-convex lenses by 2PP of the SZ2080 photoresist, a biocompatible photo-sensible material.^[33]^ Single plano-convex lenses, or a hexagonal microarray of these microlenses, were fabricated on top of a supporting borosilicate glass with a thickness of 170 μm that is integral to the design of the system. Each microlens has the geometrical parameters reported in **Table I**, and can be defined as large size micro-optics if compared with 2PP microlenses already present in the literature.^[34]^ This makes the optimization of the fabrication process challenging if we want to match a reasonable prototyping time while maintaining the structure stability and optical surface quality. To do so we devised combining surface 2PP of the outer surface of each lens (**Fig.2A**) with UV polymerization of the photoresist within the volume (**Fig.2B**)^[23,35– 37]^ implementing a higher polymerization voxel and adopting a raster scanning during 2PP (**Fig.2C**).

**Figure 2.**
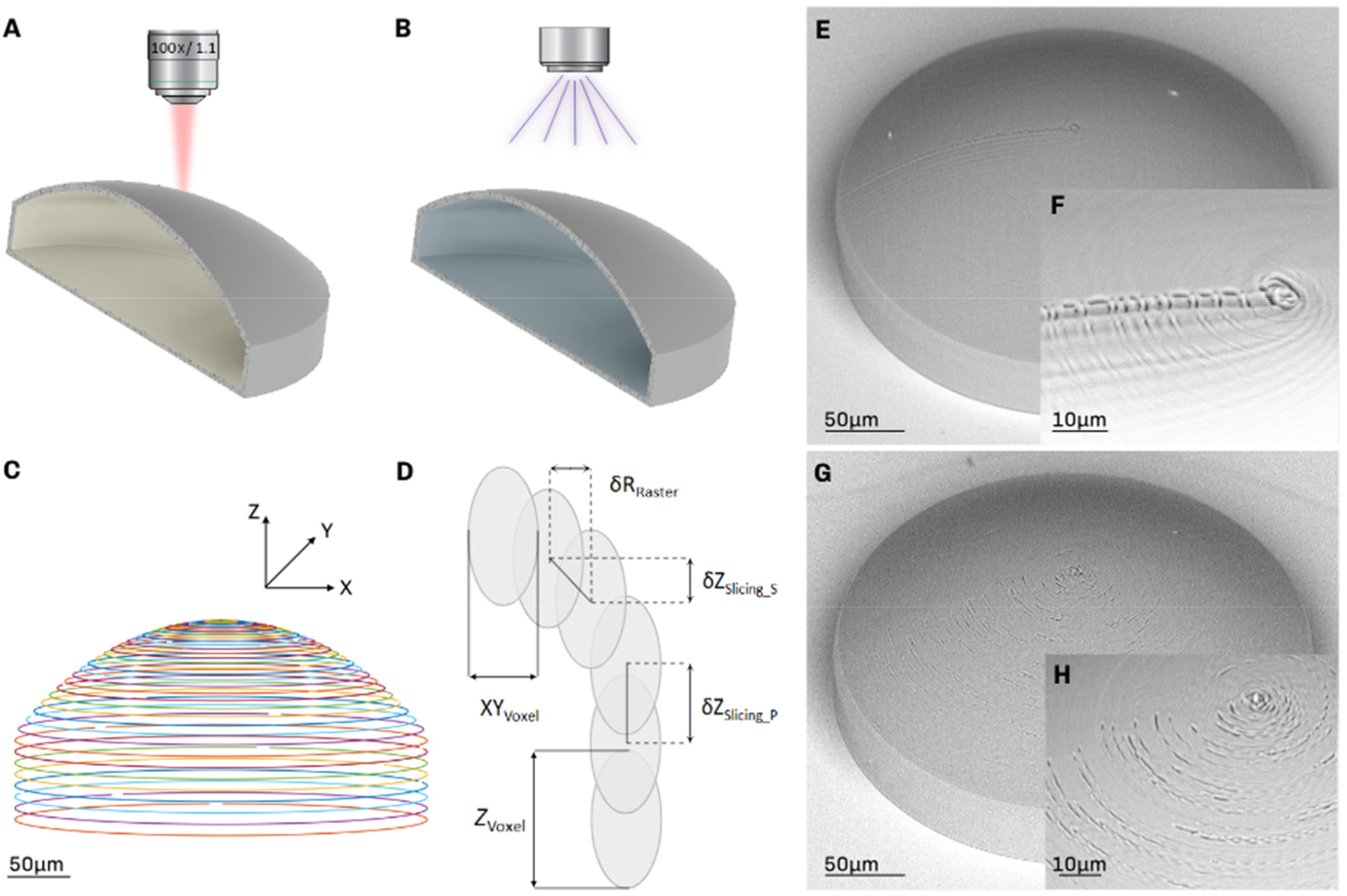
Scheme of the microlens fabrication protocol. **A**: 2PP of the lens outer surface focusing the laser beam with a 100x water immersion objective (NA 1.1). **B**: UV exposure to crosslink the unpolymerized core of the lens. **C**: rendering of the outer shell of the microfabricated lens obtained from the G-code as a sequence of concentric cycles with decreasing radius along the optical axis of the lens. **D**: details of the contour parameters of rasterization and slicing on both the pedestal (δZ_Slicing_P_) and spheric (δZ_Slicing_S_, δR__Raster_) part of the lens, and three-dimensional resolution of the polymerization voxel (XY_voxel_, Z_voxel_). **E**: SEM image of the deterministic polar scanning (non-random distribution of the start/end writing points). **F**: detail of the hinge on the outer surface due to the same start Y coordinate at each rotation. **G**: SEM image of a microlens after the randomization of the distribution of the start/end points for each rotation. **H**: detail of the overexposure effect due to the acceleration and deceleration of the translation stages during laser irradiation (random start-end point within an angle of 120 degrees).

The thickness of the 2PP polymerized surface is a crucial parameter to obtain a construct with sufficient mechanical strength to withstand the manipulation of the sample before the UV polymerization. The thickness of the shell is directly related to the 3D voxel size, which is settled by the laser focusing conditions. We decided to focus the laser beam directly into the resist using a 100x water immersion objective (see Materials and Methods section). Introducing a mismatch between the refractive index of water (1.33) and the photoresist (1.47) increases the voxel height (along laser propagation direction) due to spherical aberrations.^[38]^ Furthermore, to increase the voxel in the transversal direction we did not completely fill the objective back entrance. In this way, we obtain a voxel size of 1 μm in the xy and 5 μm in the z directions, which is a rather good compromise between the fabrication resolution and the thickness of the spherical shell.

For the polymerization of the microlens contour we adopted a raster polar scanning carried out by a concentric circular trajectory of the impinging laser beam around the lens axis (**Fig.2C**). Starting from the CAD design of the lens from Zemax calculations, we used a dedicated code to slice the model and to obtain a matrix of points in cartesian coordinates for each slicing plane. The trajectory for the microfabrication (machine G-code) has been defined as a sequence of concentric circles with a decreasing radius along the optical axis of the lens (**Fig.2C**). A key factor is to ensure a good superposition of the polymerized circles which has been obtained setting both the vertical step *δZ*_*slieing*_ (*δZ*_*slieing*_ ≤ 1 *μm*) and the radial step *δR*_*Raster*_ (*δR*_*Raster*_ ≤ 0.5 *μm*) for each vertical plane, relative to the curvature of the spherical surface (**Fig.2D**).

The fabrication process was then carefully optimized to obtain the smoothest surface for the microlenses. In the first outcomes of fabrication, the spherical outer surface was affected by a visible hinge related to the start and end points of each polymerized circle and by an overexposed region on the top of the lens due to the very dense overlap of the polymerized lines on that final region (**Fig.2E-F**). Therefore, we implemented in the code a pseudo-random distribution of the start/end writing points (Matlab function) within an angle of 120 degrees.

The homogeneity of the outer surface increased (**Fig.2G)**, despite the fact that there are still some overexposed regions, distributed all around the surface, related to the writing acceleration and deceleration during the rotations (**Fig.2H**). To avoid this effect, we added a rotation without irradiating the photoresist before and after each polymerization cycle. The quality and the smoothness of the outer surface were visibly improved, even if the central region on the top plane of the lens is still affected by the overexposure. Therefore, according to the planar and vertical resolutions of the polymerization voxel, the fabrication protocol was optimized by excluding a central area of the top of the lens with radius ∽ 1 μm from the 2PP process. Still, some faint effect of the overexposure was present (**Fig.3A**). As literature evidence demonstrates, the complete homogeneity of the central part is a common and crucial issue related to the technique and the process of fabrication.^[36]^ We envisaged that the top polymerization area could be then optimized by slowly reducing the translation speed and laser power (1-2% in each step) in the last 2PP loops.^[39]^

**Figure 3.**
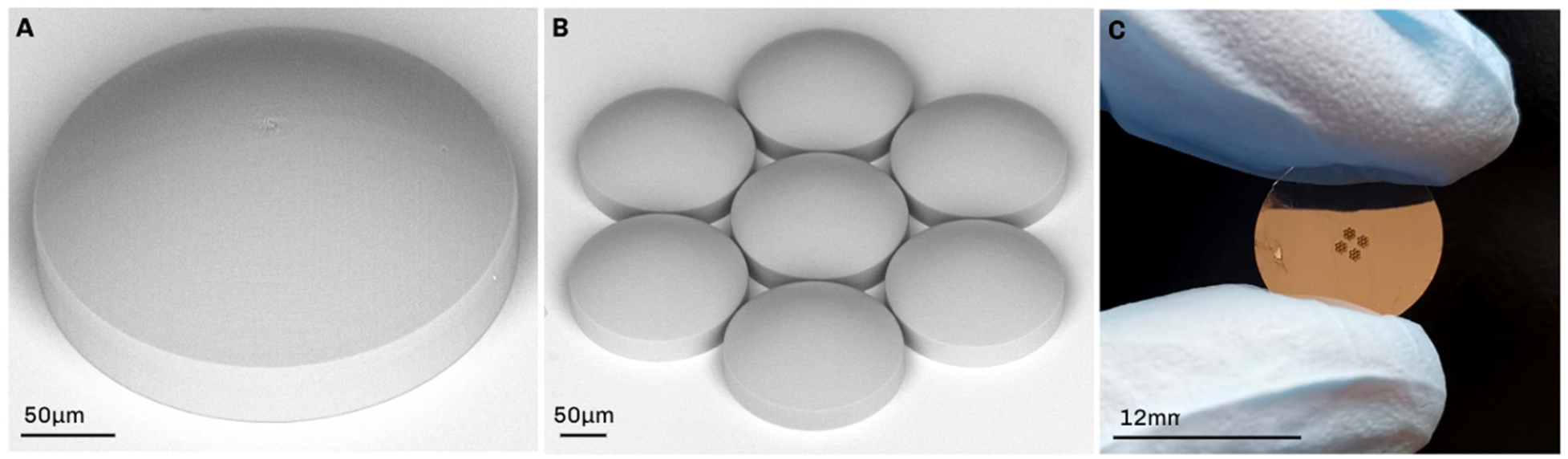
Final outcome of fabrication. **A**: SEM image of one microlens obtained from the optimization of the 2PP fabrication process and UV exposure. **B**: SEM image of one array of seven microlenses symmetrically set in a hexagonal configuration covering an area 800 μm in diameter. **C**: Image of a sample with 4 arrays of seven microlenses each covering an area 1.5 mm in side.

After the 2PP of the outer surface of the microlens, the surrounding unpolymerized photoresist is removed and then the core of the lens has to be crosslinked. For this purpose, we chose to cure the lenses by UV irradiation instead of using the more time consuming thermal approach.^[34]^ Two strategies were investigated: the direct and the indirect (through the supporting glass slide) exposure of the microlenses to the UV source. For the direct exposure case, we observed that 40 seconds of UV exposure time with UV power ranging from 100 mW to 150 mW, was not enough to completely crosslink the unpolymerized inner core of the microlens. The fabricated structures were not stable. However, the use of higher powers (up to 200 mW) and/or the increase of the exposure time till 120 seconds (step of 40 sec) promoted the degradation of the resist, for instance the external surface already polymerized tends to become stiffer turning yellow (**SI10.C**). With the indirect exposure approach, we can instead identify optimal processing conditions for the crosslinking of the inner core of the microlenses (see **Fig. SI10.D, E, F**). Low UV powers (≃ 200 *mW*) were not enough to crosslink the overall volume of resist within the construct, causing cracks in the lens base and the collapse of the lens (See **Fig. SI10.D**). On the other hand, the use of higher UV powers (≥ 440 mW) induced the distention of the outer surface improving its smoothness, but it opened a hole in the central part of the dome due to the excessive expansion of the inner resist caused by the use of too high UV power (see **Fig. SI10.F**). So, we identified 320 mW as the best and suitable value which allowed us to achieve the complete crosslinking of the microlens volume without compromising the stability and the surface of the structure (see **Fig. SI10.E**).

The inspection of the microlens surface with a tip profilometer demonstrates that the lens surface profile is spherical with a radius of curvature of *R* = 193 ± 4 *μm*, therefore only 4% smaller than the design (see **SI 6, “Profilometer analysis of the shape of the microlenses”**). This slight deviation of the curvature radius in the fabricated surface, from the desired one, is expected due to the dimensions of the polymerized voxel and some possible shrinking occurring upon developing the microstructure. If needed, this negligible deviation can be corrected by working on the CAD shape of the lens but for our application is negligible as will be demonstrated in the next section. Moreover, with the perspective of fabricating micro-lenses with an even bigger volume, which might collapse for example in the pedestal section, the stability can be further improved by increasing the thickness of the 2PP shell by a multi-scan approach.

In summary, the optimized fabrication protocol, based on a combination of 2PP and UV induced polymerization, promotes a 98% reduction of the fabrication time, if compared to the one required to 2PP the whole volume of a full lens (from 12 hours to 8 minutes) without compromising the stability and the homogeneity of the microlenses. At the same time, the appropriate choice of the slicing (*δZ*_*slieing*_ = 1 *μm*) and radial (*δR*_*Raster*_ = 0.5 *μm*) steps within the 2PP fabrication phase of the protocol, allowed us to obtain a lens surface quality appropriate to perform optical imaging (as better shown in the optical characterization section) and with a good reproducibility (±5%) of the shape (**see SI6, “Profilometer analysis of the shape of the microlenses”**). In addition, an even smoother surface can be obtained, by decreasing the imposed radial step to *δR*_*Raster*_ ≃ 0.3 *μm*, during the polymerization at the expense of the fabrication time. Fabrication time increases from 8 minutes per lens at *δR*_*Raster*_ ≃ 0.5 *μm* to 15 minutes per lens at *δR*_*Raster*_ ≃ 0.3 *μm*. Overall, we obtain stable microlenses with optical quality and with a higher size if compared with the microlenses fabricated by 2PP in the literature.^[23,25,34–36,40,41]^

The imaging area can be even increased with respect to the predicted ≃ 250 μm for a single microlens (**SI5, “Field of view of microlenses”**) by rescaling the microfabrication process to a hexagonal array of microlenses covering an area of ≃ 800 *μm* (**Fig.3B**). Four hexagonal arrays (**Fig.3C**) were fabricated on a 12mm Ø circular glass substrate (170 *μm* thickness) covering an area 1.5 mm in size. The production of one set of arrays requires less than 4 hours.

### 2.3 Optical characterization

Before assessing the possibility to get fluorescence images of cells through the fabricated microlenses, we tested their optical quality by the study of the intensity distribution of a beam propagating along the optical axis, the wavefront measurement at the focal plane and the measurement of the modulation transfer function. For all these tests, we used a collimated diode laser beam at *λ* = 635 *nm*.

#### Minimum spot size under collimated beam

The intensity distribution on planes perpendicular to the optical axis provides a direct visualization of the effect of the aberrations, each having a peculiar signature.^[42]^ The microlenses were tested in overfilling conditions to enhance the aberration effect. The intensity distributions sampled at 10 *μm* steps (**Fig.4A** and **SI7, “Intensity distribution sampling along the optical axis”**) in a range of about ±20 *μm* around the effective focal plane of the microlens (corresponding to the minimum spot size) are reported in **Fig. 4B**. These images indicate the presence of spherical aberrations as concentric circles, turning more evident for distances *z* < *f*_*FFL*_: for planes at distances larger than the front focal length, *z* > *f*_*FFL*_, as expected,^[42]^ the central peak of the intensity distributions becomes more diffuse and the concentric rings cannot be singled out. The simulated intensity distributions (**Fig.4C**) agree with the experimental ones. Due to the presence of the spherical aberration rings, we cannot define a Gaussian width (*1/e*^2^) for the intensity distribution across the optical axis (see **SI7, “Intensity distribution sampling along the optical axis”**).^[43]^ However, the full width at half maximum (FWHM) of the central spot in planes close to the plane of minimum confusion can be measured on cross-section profiles of the intensity distributions. The full width, FWHM = 0.98 ± 0.10 *μm* (averaged over an entire array of 7 microlenses), is in very good agreement with the simulation of the beam profile (**Fig.4D**). This value corresponds to a beam waist (*1/e*^*2*^ radius) *w*_0_ = 0.84 ± 0.09 *μm*. An estimate of the Rayleigh range, *z*_*R*_, can be obtained by the relation *z*_*R*_ = *πw*^4^_0_ /*λ* ≃ 2.5 ± 1.1 *μm*.

**Figure 4.**
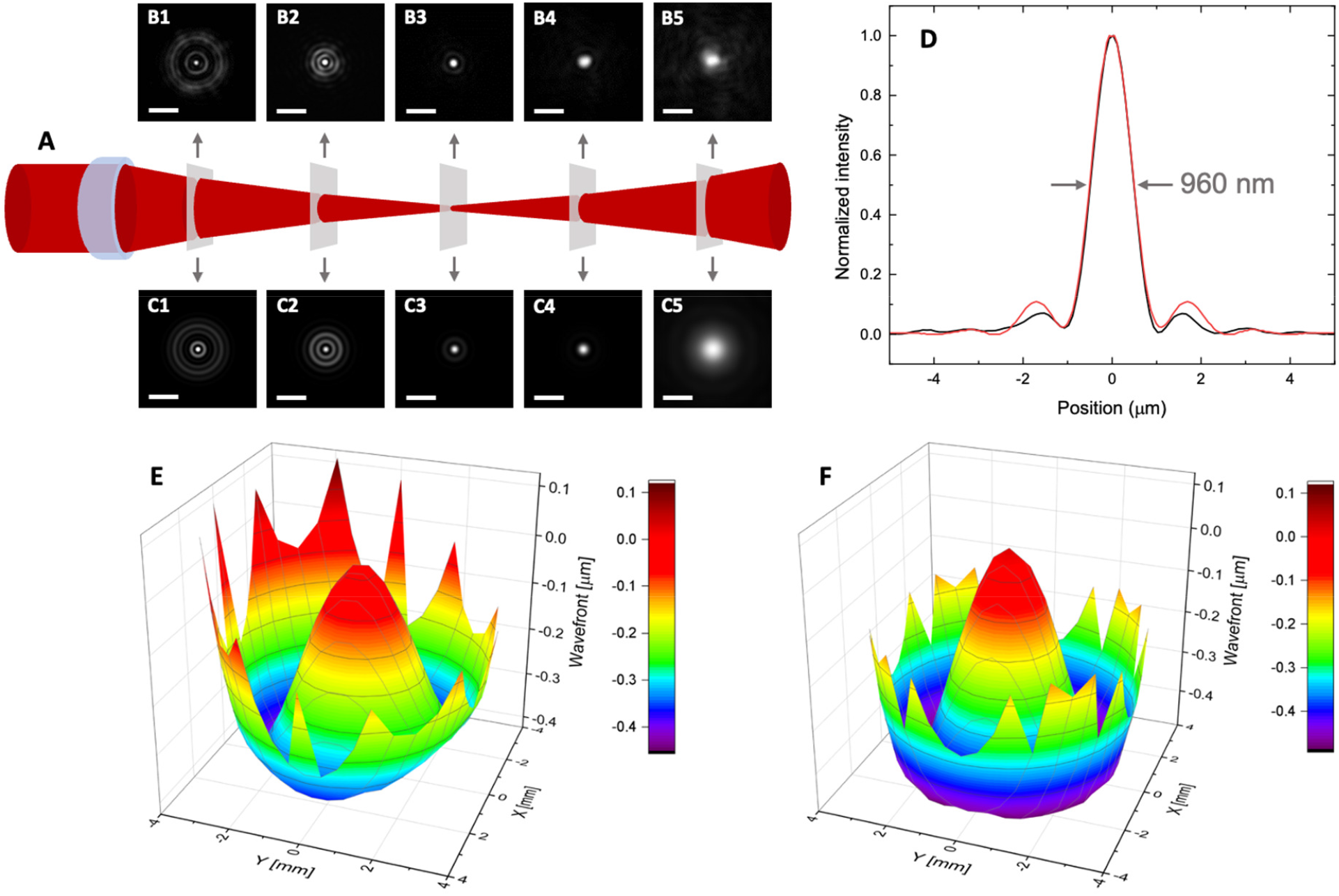
Aberrations of the microlenses. **A**: Schematics of the configuration adopted for the measurement of the intensity distribution of a Gaussian beam focused by a plano-convex microlens (**Table I**). The tube lens and the camera sensor are aligned to accept rays from infinite (see **Fig.SI7** for details). **B**: Images of the intensity profiles collected at various distances along the optical axis. The mutual distance between the planes is 10 μm, starting from the plane closer to the microlens (from left to right). Scale bars 5 μm. **C**: Simulated intensity distributions with the same 10-μm spacing along the optical axis. Scale bars 5 μm. **D**: Exemplary profile of a Gaussian beam focused by one microlens (the beam overfills the microlens). Experimental and simulated data are reported as black and red lines, respectively. The FWHM of the experimental profile is *FWHM* ≃ 960 *nm*. **E-F**: Measured (**E**) and simulated (**F**) wavefront at the focal plane (minimum confusion plane) of the microlens. Piston, tip and tilt aberrations were subtracted from the experimental wavefront for visualization.

#### Wavefront measurement

The wavefront was measured by means of a Shack-Hartman sensor (**SI4**, “**Microlens wavefront measurement**”). The resulting wavefront of the microlens, at the maximum pupil of about 264 μm is reported in **Fig.4E**. The mexican-hat shape is typical of the *Z*_4_^0^ Zernike polynomial that characterizes the first-order spherical aberration. In fact, the Zernike decomposition of the experimental wavefront gives an amplitude of the spherical component *a*_4,0_ = 0.10 ± 0.01 *μm* (averaged over an entire array of microlenses), whereas all the other Zernike coefficients are lower than 0.02 μm. The wavefront predicted for the microlens by Zemax simulations, reported in **Fig. 4F**, agrees in shape with the experimental one. The predicted value a_4,0_^(*Z*)^ = 0.15 *μm* is larger than the measured value *a*_4,0_. Indeed, while the simulated distribution of the Zernike weights is peaked on a specific aberration component (here, the spherical one), the experimental wavefront is described by a distribution of Zernike polynomials: together with the dominant spherical component, *a*_4,0_, other aberrations contribute with small, but non-zero, coefficients. As a consequence of the superposition of multiple aberration coefficients, the fit of the resulting experimental wavefront to the Zernike polynomials results in an effectively higher uncertainty on *a*_4,0_. This is probably the origin of the discrepancy between the numerical values of *a*_4,0_^(*Z*)^ and *a*_4,0_.

Still, the peak-to-valley (or Root-Mean-Square, RMS) wavefront error determines the overall optical quality more prominently than the amplitude of individual Zernike coefficients. Indeed, the average peak-to-valley wavefront distortion, after subtraction of the piston, tip and tilt components, is about *0*.*62 ± 0*.*04 μm* and it is similar to the simulated value (**Fig.4F**) of 0.51 *μm*. Analogously, the average RMS value is 0.13 ± 0.02 *μm* and is in good agreement with the Zemax prediction, RMS = 0.15 *μm*.

Overall, the *a*_4,0_ and RMS wavefront error of the microlens reveal an even better behavior of the microlens when compared to a commercial macroscopic plano-convex lens (f = 25mm, Thorlabs, LA1252) with ∽35% pupil size (see **SI4, “Microlens wavefront measurement”**).

#### Modulation transfer function

The MTF can be measured pointwise by means of the analysis of a reference target, such as the USAF 1951. A qualitative result of the imaging of the USAF target through the microlenses is given in **Fig.5A-B**. The microlenses are used in a finite conjugate condition (**Fig.5H** and **Fig.SI3.1A**), with a magnification determined by the ratio of the distance (q) between the image plane and the exit principal plane over the distance (p) between the sample plane and the entrance principal plane. A 20x microscope objective collects the image produced by the microlens in the objective focal plane.

**Figure 5.**
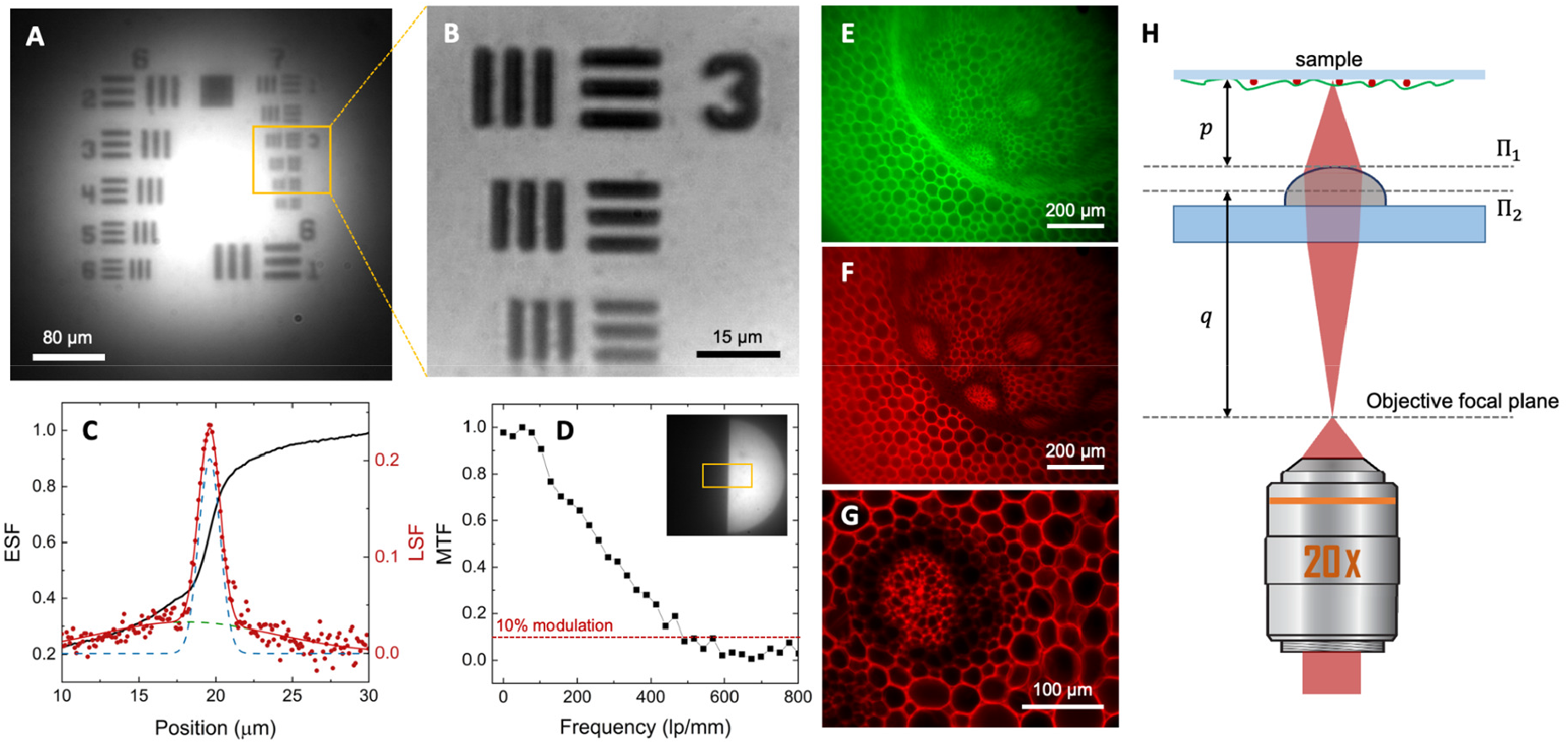
Optical performance of the microlenses. **A-B**: Example of wide-field images of an USAF1951 resolution test chart obtained by a single microlens and a 20x objective as described in the text. Depending on the adopted relative positions of the microlens and the objective the overall magnification equals 6x (**A**) and 30x (**B**). **C**-**D**: Measurement of the MTF (Modulation Transfer Function) of the microlenses by the knife edge method. **C**: Edge Spread Function (ESF, black solid line) and Line Spread Function (LSF, red circles) of a single microlens. In order to eliminate the background contribution, the LSF is fitted to a two-component Gaussian function (dashed lines), the background wider Gaussian component is subtracted, and the resulting LSF is Fourier transformed to retrieve the MTF. **D**: MTF of the microlens derived as the Fourier transform of the background-subtracted LSF in panel **C**. Inset: Image of a sharp razor blade produced by the microlens and exploited for the ESF quantification. The ESF is extracted as the average intensity profile along the horizontal axis within the yellow Region of Interest; the LSF is computed as the first derivative of the ESF. **E-F**: Wide-field images of a sample of Convallaria Majalis obtained under blue (E) and green (F) excitation and epifluorescence collection. **G:** Control image of the same sample obtained on a wide-field microscope (Materials and Methods section). **H**: sketch of the finite conjugate use of the microlens. *п*_1_ and *п*_2_ are the two principal planes.

The MTF of the fabricated microlenses was derived by inserting a sharp edge in the sample plane, and by collecting and digitizing an image to retrieve the Edge Spread Function (ESF) (**Fig.5C,D**). We computed the derivative of the ESF to obtain the Line Spread Function (LSF), whose Fourier transform is the Modulation Transfer Function (MTF, **Fig.5D**). The MTF, that is the modulus of the Fourier Transform of the point spread function, offers a complete evaluation of the optical response of the microlenses for imaging purposes.^[44]^ A 10% modulation is reached at a cutoff frequency of about 490 lines pairs/mm, which is comparable to the cutoff frequency of more sophisticated microfabricated optics in the recent literature.^[25]^

The optical response of the microlenses can also be characterized by an interferometric measurement of the lens phase profile as recently implemented by Zhao et al.^[45]^ for metalenses. By inserting the metalenses in one of the two paths of a Mach-Zender interferometer (**Fig.SI9.1**) one can measure the full phase profile of the metalenses. For a refractive lens, we need to compute the phase shift induced by refraction, taking into account the shape of the curved surface, its thickness and the index of refraction as we do here in section **SI9**, “**On-axis holographic analysis of the profile of the microlenses**”. By adopting an on-axis holographic measurement we are able to sample the phase profile induced by the microlens over an optical path difference larger than 40 × 2*π*. We have compared the measurements with the predictions made for a perfect spherical shape of the microlens (**Fig.SI9.2C**, characterized by the sag function, 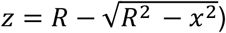 and with those made for its parabolic approximation, 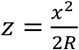 (**Fig.SI9.2-C**). We find that both predictions agree well with the measurements when the curvature radius is *R* = 195 *μm*, with a slight but definite improvement for the parabolic shape (see **Fig.SI9.2-D)**. The value of the radius found here agrees very well with the evaluation done by means of the profilometer analysis (*R* = 193 *μm* ± 4 *μm*).

Having well characterized the on-axis optical properties of the microlenses and measured directly the microlens profile through the phase of the field at its exit pupil (see **SI9, “On-axis holographic analysis of the profile of the microlenses**”), we can simulate the off-axis behavior of the microlens. We are coupling the microlenses to a microscope objective, which is a sample space telecentric lens as depicted in **Fig.SI3.2A-B**. As reported in **Fig.SI3.2C-F**, the off-axis optical behavior of the microlens indicates the possibility to use it for raster scanning optical imaging. In fact, over a field of view (on the sample) of about 64 *μm*, we find that the RMS spot radius changes from 280 *nm* (on axis) to 360 *nm* at a field of view of 50 *μm*. It then increases rapidly to 1.6 *μm* at the field of view edge of 64 *μm*. This resolution value is compatible with the extent of the MTF up to 600 lines/mm (see **Fig.5D**). The off-axis optical behavior of the fabricated microlenses indicates again the possibility to use them for raster-scanning optical imaging.

#### Cell imaging through the microlenses

As a proof of concept, microlenses can be coupled to a wide-field microscope to collect wide-field images both in transmission and in fluorescence mode, or can be coupled in a virtual image configuration to a scanning optical microscope (see **SI3, “Microlens coupled to a raster scanning optical microscope”**). In the first case, the microlenses are used in the same finite conjugate condition (**Fig.5H** and **SI3**) exploited for the USAF target imaging in **Fig.5**. Examples of fluorescence wide-field images collected in this configuration are reported in **Fig.5E,F** for a sample of Convallaria Majalis, excited in the blue (**Fig.5E**) or the green (**Fig.5F**) spectral bands. From these images, very similar to the ones obtained using a wide-field microscope **(Fig.5G**), we can clearly distinguish the most relevant parts of the root. The pericycle and endodermis appear as a bright green circle in **Fig. 5E**, surrounding a set of xylems and phloems and surrounded by the cortex. A slight field curvature can be observed at the edge of the field of view in **Figs.5E** and **5F**, compared to the reference image in **Fig.5G**.

For non-linear excitation microscopy it is essential to couple the microlens with a scanning setup. In this second type of application, we couple the microlenses directly to a low numerical aperture objective in a raster scanning optical microscope. The microlens is set at a distance from the exit lens of the objective smaller than the objective working distance (see **Fig.6J** and **SI3, “Microlenses coupled to a raster scanning optical microscope”**). In this configuration the microlens creates, for any position of the laser beam during the scanning, a virtual image of the sample that is then collected by the microscope objective. The magnification of the resulting image can be computed as the product of the magnification of the objective and that of the microlens. The latter depends directly on the distance, *δz*, between the microlens and the sample plane of the microscope objective (for details see **SI3**, “**Microlenses coupled to a raster scanning optical microscope**”). This configuration can be implemented both on confocal and on non-linear excitation fluorescence microscopes, also in the case of a microlens array, as reported in **Fig.6**. In the example of **Fig.6**, the sample is a cell culture (human dermal fibroblasts) grown on the very same glass slide (opposite side) where the microlenses are fabricated. Fibroblasts can be imaged by the microlenses by properly adjusting the position of the microlens array with respect to the microscope objective. When *δz*≃ 0, only the cells not covered by the microlenses get imaged, with the microscope objective magnification (**Fig.6A** and **Fig.6D**) on the confocal microscope. When *δz* ≃165 *μm*, one can get images of the cells through the microlenses (**Fig.6B,C**). The estimated transverse magnification due to the microlens (see **SI3**, “**Microlenses coupled to a raster scanning optical microscope**”) is *M*_*T*_ ≃ 2.4. 3D reconstruction of the cell culture is possible too, as shown in **Fig.6E**.

**Figure 6.**
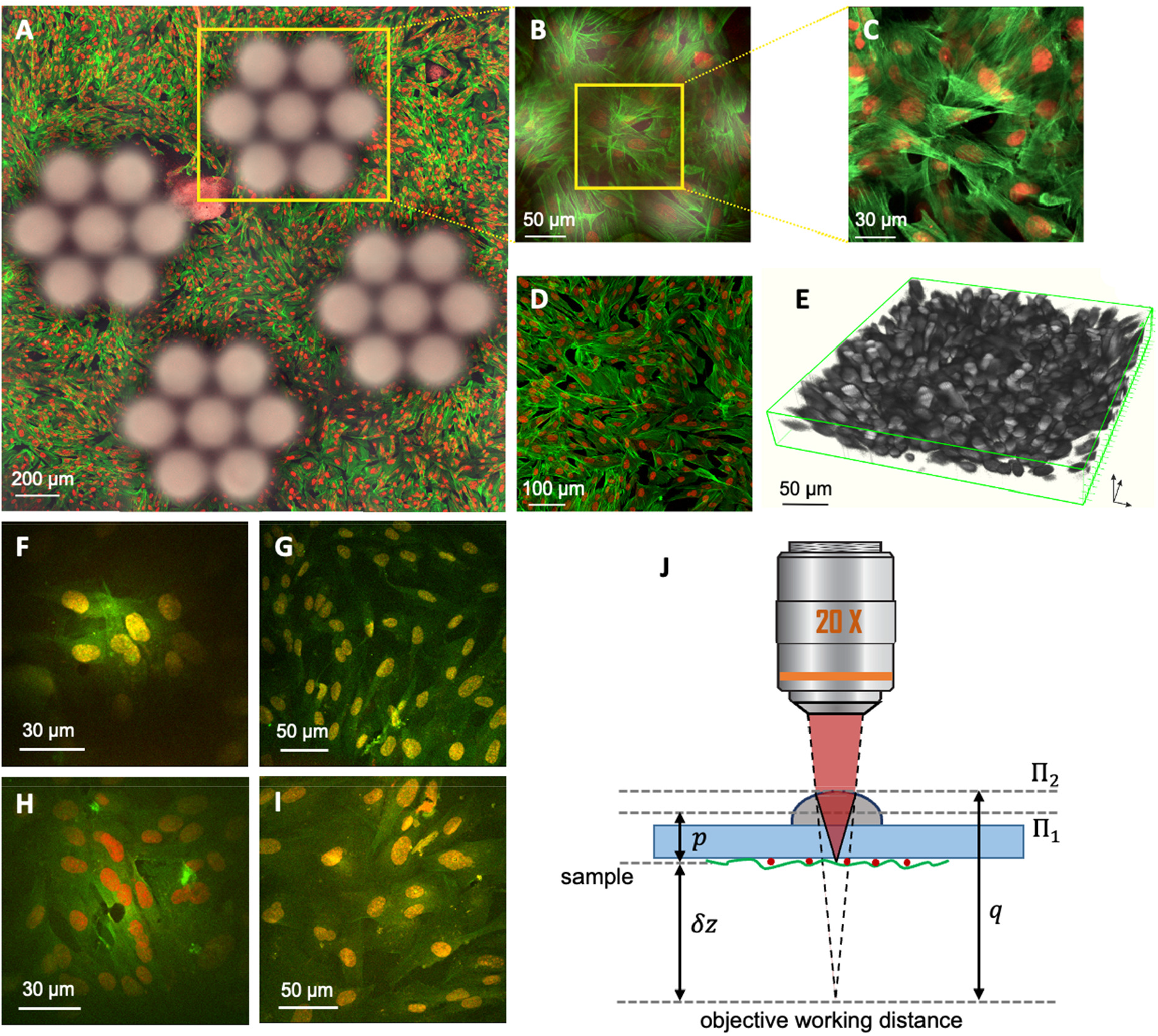
A: Confocal image (3×3 stitched tiles) of stained human fibroblasts (FITC-phallodin and DRAQ5/Hoechst staining). The fluorescence confocal and transmitted images at the glass coverslip focal plane are superimposed. Microlenses are imaged through out-of-focus transmitted laser light. Cell nuclei, in red, and cytoskeleton, in green, are visualized in fluorescence (actin, excitation at 488 nm, emission at 525/50 nm; nuclei, excitation at 647 nm, emission at 665/81 nm). **B**: Full field-of-view fluorescence confocal image through an array of microlenses, superimposed to the transmitted-light image. The cell fluorescence image is collected when the sample is shifted by *δz* = 165 *μm* with respect to the objective focal plane. **C**: Cropped scan area through a single microlens. **D**: Full field-of-view confocal microscope image of cells at the glass coverslip focal plane (*δz*≃0, scale bar 100*μm*). A 20x dry objective was employed for **A-D. E**: 3D reconstruction of cells through an array of microlenses. The image was rescaled accordingly to the microlenses magnification: *M*_*T*_≃ 2.4, *M*_*z*_ ≃*M*_*T*_^2^. **F-I:** Fluorescence images of the cells under two-photon excitation (*λ* = 800 *nm*). **F**: Fluorescence image collected through the microlenses coupled to a 20x dry objective at a distance *δz* = 135 *μm* (with respect to the objective sample plane), resulting in a total magnification *M*_*tot*_ ≃ 45. **G**: Control image obtained through the 20x dry objective at *δz*≃0. **H**: Image of the cells through the microlenses coupled to a 25x water immersion objective obtained at *δz* ≃43 *μm*. **I:** Control image obtained with the 25x water immersion objective when *δz*≃0. **J**: Sketch of the configuration used to couple the microlens with the optical scanning microscope. The distance between the sample (dark gray) and its virtual image (light gray) through the microlens is *δz*. П_1_ and *П*_2_ are the two principal planes of the microlens.

The microlenses optical quality is sufficient to induce also two-photon excitation and to collect non-linear excitation fluorescence images both when used in air (**Fig.6F**) and when used with water immersion objectives (**Fig.6H**), despite the lower dioptric power of the microlenses, due to partial index matching. When the microlenses are coupled with a 20X dry microscope objective we found that the images of the cells through the microlenses can be recovered at *δz* = 135 *μm* resulting in a magnification ≃2.3, that leads to a total magnification *M*_*tot*_ = *M*_*T*_ *M*_*obj*_ ≃2.3 × 20≃46. When the microlenses are coupled to a 25X water immersion objective, the images of the cells through the microlenses can be recovered at *δz* = 43 *μm*, corresponding to a microlens magnification *M*_*T*_≃1.75, that leads to a total magnification *M*_*tot*_ = *M*_*T*_ *M*_*obj*_ ≃1.75 × 25≃44. Indeed the raster scanning images obtained with the two microscope objectives are quite similar. The field of view (FOV) is reduced by a three/fourfold factor, with respect to the FOV of the objectives, depending on the coupling distance between the microlenses and the objective.

Notably, the perceptive quality, the resolution and the signal to noise ratio of the images collected through the microlenses are very similar to the ones obtained on the raster scanning optical microscope in its conventional configuration. The application of the “Perception based Image Quality Evaluator” (PIQE),^[46]^ one of the available “*No-Reference Image Quality Assessment”* algorithms, to the images reported in **Fig.6** gives the results summarized in **Table 2**. Even though there is a substantial loss of perceptive quality when microlenses are used on the confocal setup, the quality of the TPE images is not substantially affected by the use of the microlenses.

**Table 2.**
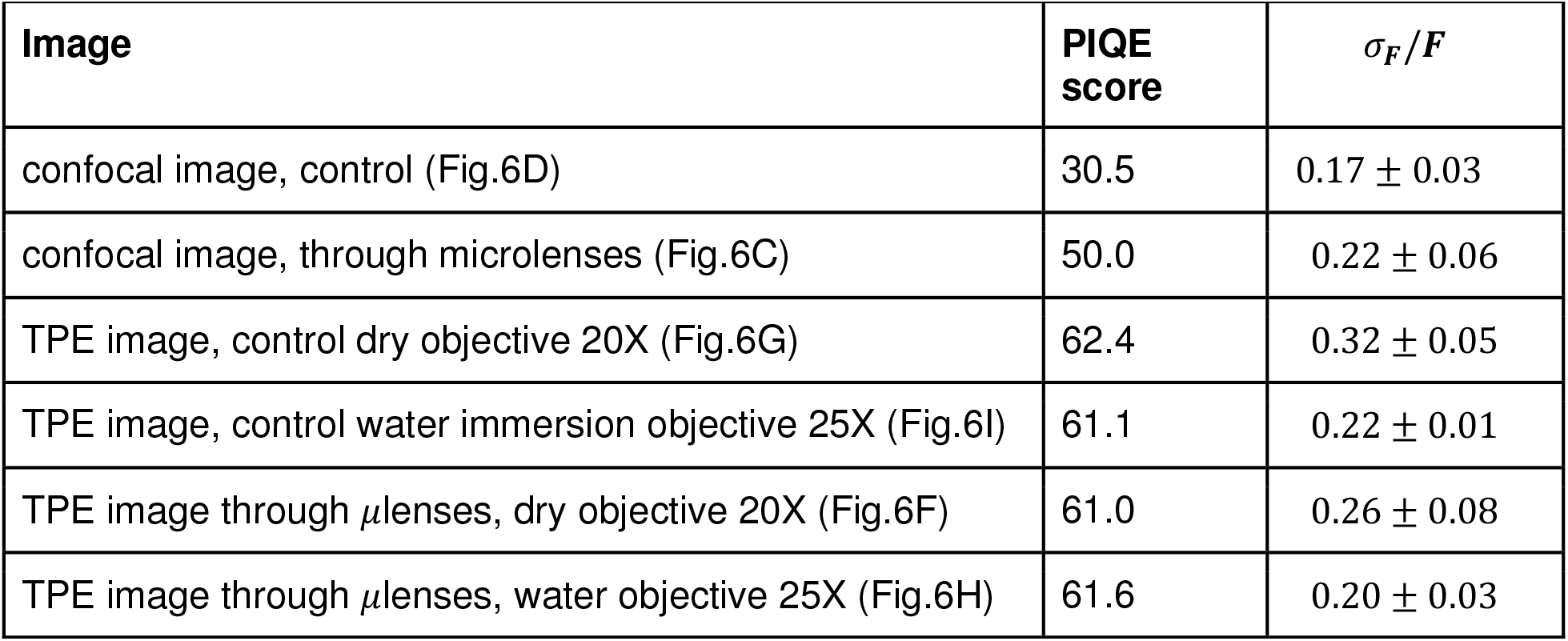
evaluation of signal/noise metrics on the confocal and non-linear excitation images. Application of the PIQE evaluator^[46]^ to images taken through the confocal or the TPE microscopes, with or without the coupling to the microlenses. The best quality image is assigned a PIQE = 0.

| **Table 2: evaluation of signal/noise metrics on the confocal and non-linear excitation images.** |  |  |
| --- | --- | --- |
| **Image** | **PIQE score** | **$\sigma_F/F$** |
| confocal image, control (Fig.6D) | 30.5 | $0.17 \pm 0.03$ |
| confocal image, through microlenses (Fig.6C) | 50.0 | $0.22 \pm 0.06$ |
| TPE image, control dry objective 20X (Fig.6G) | 62.4 | $0.32 \pm 0.05$ |
| TPE image, control water immersion objective 25X (Fig.6I) | 61.1 | $0.22 \pm 0.01$ |
| TPE image through μlenses, dry objective 20X (Fig.6F) | 61.0 | $0.26 \pm 0.08$ |
| TPE image through μlenses, water objective 25X (Fig.6H) | 61.6 | $0.20 \pm 0.03$ |

As a second image quality estimator, the ratio *σ*_*F*_/< *F* > of the standard deviation, *σ*_*F*_, to the average fluorescence signal *< F >* was evaluated on 4 to 7 ROIs of identical size (22×22 pixels) on regions of uniform signal signal. If the microlenses collected the light with an overall lower efficiency, we should obtain larger values of *σ*_*F*_/< *F* >. The values of this parameter (see **Table 2** and **Fig. SI8.1**) indicate again that the relative noise on confocal images is not increasing substantially (within the standard deviation) when coupling microlenses to the microscope objective.

Also in terms of spatial resolution, the use of microlenses coupled to the raster scanning microscope, does not induce any dramatic loss. We estimated the effective spatial resolution by measuring the minimum size of the features on the images. For the confocal images, we analyzed the thickness of the actin filaments (**SI8, “Analysis of the noise on the two-photon excitation images through microlenses**” and **Fig. SI8.2**) finding that the average size of the features (likely thin bundles of actin filaments) are *δx* = 0.8 ± 0.2 *μm* and *δx* = 1.0 ± 0.3 *μm* for images taken respectively on the confocal microscope and through microlenses coupled to the confocal microscope.

For the case of TPE images, the size of the chromatin inhomogeneities in the cells nuclei was considered, finding a radial *1/e*^2^ radius < *δr* > = 2.3 ± 0.6 *μm* and < *δr* > = 2.0 ± 0.6 *μm* for the TPE microscope (water immersion objective) and for the case in which the microlenses were coupled to the same TPE microscope objective (see **Fig. SI8.3** for details). Both for confocal and TPE microscopy, we do not find any evidence of substantial degradation of the spatial resolution. However, the analysis given above does not substitute a direct measurement of the spatial resolution on sub-resolved objects.

In this report, we do not apply microlenses directly to the imaging of thick specimens. However, the analysis reported above allows us to draw a few general practical considerations on the use of the microlenses in optical microscopy of tissues in-vivo. The coupling of microlenses to the objective of a scanning microscope implies a calibration step. In fact, the actual position of the sample plane, the magnification and the field of view all depend (**Fig. SI3.3**) on the distance between the objective front lens and the microlens entrance principal plane. For the microlens fabricated here, the position of the sample plane on which the raster scanning is actuated changes from about 5 *μm* to about 44 *μm* and the magnification changes in the range 3 < M < 9 (microlens focal length *f*_*μ*_ = 240 *μm*). This range reduces to 2.1 < M < 7 when *f*_*μ*_ = 340 *μm* (**Fig.SI3.3**). A consequence of this behavior is the need to have a reference for the size in the focal plane at the various slices of a Z-scan. This can be obtained, for example, by fabricating on the sample space a reference cone, as done for example in ref. [11].

## Conclusions

In summary, we have shown that microlenses, with high numerical aperture (0.4) and large diameter (260 *μm*), can be fabricated with a medium throughput, about 8 minutes/lens, by limiting the 2PP fabrication to a ≃ 1 *μm* thick outer crust and a post-development volume polymerization. This procedure leads to a 98% reduction in the fabrication time with respect to volume polymerization, still obtaining sufficient optical quality to use the lenses for direct wide-field imaging of cells, both in transmission and in fluorescence mode. No evidence of inhomogeneities in the lens volume is found due to the post-development processing.

Wide-field imaging through the microlenses can be obtained in a finite conjugate coupling approach, where the microscope objective and the microlenses are arranged in series. This method can be used for routine testing of the optical quality of the microlenses. More interesting, the microlenses can be coupled to a raster scanning non-linear excitation optical microscope, even by using low numerical aperture and low magnification objectives: in this case a virtual image configuration is adopted. Our experiments (**Fig.6**) show for the first time that it is possible to use the microfabricated lenses to efficiently induce non-linear excitation and collect two-photon excitation images from cells with the same level of laser intensity and similar signal to noise ratio. This work opens the possibility to implement micro-optical devices for non-linear excitation that, due to the limited size, could be used as implants for the direct and continuous optical inspection of biological dynamics *in vivo*.

## MATERIALS and METHODS

### Sample Preparation for Two-Photon Laser Polymerization

The photoresist used for the 2PP of the microlenses was a negative hybrid organic-inorganic UV-curable resist known as SZ2080TM,^[47]^ biocompatible and validated for biological applications.^[11,48]^ To increase the efficiency of the polymerization 1% of Irgacure-369 (2-Benzyl-2-dimethylamino-1-(4-morpholinophenyl)-butanone-1) is added to the resist as photoinitiator. About 46 μl of liquid SZ2080 photoresist were deposited by drop casting on a 12 mm diameter circular glass coverslip with a thickness between 170 and 230 μm (#1.5, Menzel-Glaser, Germany). To allow the drying of the resist thus to prevent any unwanted sliding of the structures during the fabrication process, the samples were left under the chemical hood for 48 hours. Once the material had condensed, the photoresist reached a semi-solid state suitable for the further laser-induced polymerization.

### Two-Photon Laser Polymerization Setup

The 2PP fabrication was performed by a femtosecond Ytterbium (Yb) fiber laser (Satsuma, Amplitude System). The laser wavelength was 1030 nm, the pulse duration is ∽280 fs and the repetition rate was fixed to 1MHz. The output power reaches a maximum value of 10 W. The laser beam passed through a series of mirrors that steered it to a 2x telescope and a half-wave plate coupled to a polarizer (MPS50GR-TTM-G80-DC-LMO-PLOTS, Aerotech, USA) which was used as a system for the power control. The half-wave plate was mounted on a software controlled motorized rotator which allows the dynamic control of the power during 2PP. At the end of the optical path the beam was tightly focused by a water matched CFI plan 100XC W objective with a numerical aperture (NA) 1.1 (Nikon, Japan) directly onto the photosensitive resist. The sample was mounted within a glass container with a diameter of 100 mm connected to a gimbal mechanical system (Gimbal Mounts 100, Thorlabs). The sample holder-gimbal complex was screwed on a planar (X, Y) translational stage (ANT95-50-XY, Aerotech, USA) while the motion of the objective along the Z-direction was controlled by a high resolution linear stage (ANT130-035-L-ZS, Aerotech, USA) balanced by two air compressed pneumatic pistons. The motion of this three-axes stages was controlled by software (Automation 3200 CNC Operator Interface, Aerotech, USA) as well as the half-wave plate rotation and they were provided with a feedback position and velocity control system with a resolution on the order of 200 nm. For a real time monitoring of the writing process within the working area, as well as the polymerization process, a red-light emitting diode illumination was positioned in the empty central cavity of the gimbal, under the sample holder-gimbal complex, and a CCD camera was mounted behind a dichroic mirror (Thorlabs, Germany). The three translational stages and all the components of the optical path were placed upon a granite table (Zali Precision Granite Technology, Italy).

The outer surface of each microlens was two-photon polymerized directly focusing the laser beam into the SZ2080 photoresist droplet, moving the sample at a speed of 1 mm/s. Due to laser radiation absorption, the pulse energy used for 2PP of the microlenses depends on the SZ2080 droplet thickness thus ranging from 13 nJ to 15 nJ (at the objective entrance pupil).

### Sample Development and UV Exposure

After the two-photon polymerization process, the samples underwent a development procedure to remove the unpolymerized photoresist surrounding the volume of the microfabricated microlenses. The samples were soaked in a glass beaker filled with a 50% v/v 2-pentanone, 50% v/v isopropyl alcohol solution (Sigma-Aldrich, USA). After 30 min, they were cautiously washed with a few drops of isopropyl alcohol then gently dried by room temperature Nitrogen. Afterwards, the sample was UV irradiated passing through the glass substrate in order to avoid the direct and additional exposure of the previously polymerized surface. Consequently thus the radiation intensity and the time of exposure depend on the thickness of the substrate, the volume of photoresist to be polymerized and the shape of the lens. Therefore, the process was optimized exposing the sample to 300 mW of UV irradiation (λ=385nm) for 120 seconds (LIGHTNINGCURE LC-L1V3) in order to crosslink the unpolymerized core of the lens achieving the structural stability without causing resin degradation.

### Optical and Scanning Electron Microscopy Qualitative Characterization

Qualitative analysis of fabrication outcomes in terms of shape and mechanical stability were performed by an optical microscope with different magnitudes 4×, 10×, 20×, 40× (Eclipse ME600, Nikon, Japan) and by a scanning electron microscope (SEM, Phenom Pro, Phenom World, Netherlands). The SEM observations were carried out at 10 kV.

### Profilometry Analysis

The analysis of the overall shape of the lenses were performed by a surface profilometer (KLA Tencor P-17) using a 5 um stylus tip radius with cone angle 60 degrees and setting the scan speed as 20 μm/s, 20 Hz sampling rate, 1 mg as the down force to be applied on the sample surface (see **SI6, “Profilometer analysis of the shape of the microlenses”**, and **Fig.SI6**). The scanning of each lens was performed by sampling it each 5 μm along both the planar directions (X,Y) and both the positive and negative sense, within a scanning box of 350 μm side. Lastly, the surface profile has been interpolated with a spheric function in order to evaluate the reliability of the fabrication with respect to the lens model.

### Fluorescence Microscopy

For confocal imaging human dermal RFP fibroblasts (Innoprot, Spain) were cultured in Dulbecco’s modified Eagle’s medium (DMEM) supplemented with 10% fetal bovine serum, 1% L-glutamine (2 mM), penicillin (10 units/ml),and streptomycin (10 μg/ml) at 37°C and 5% CO_2_. Then, 20k/50μl cells were drop-seeded on borosilicate glass coverslips,and after 4 hours of incubation, diluted in 3 ml of DMEM and expanded for 48 hours. Then, cell culture was fixed in 4% paraformaldehyde (Sigma Aldrich, Italy) for 5 minutes and stained, for nuclei identification, with: DRAQ5 (ThermoFisher Scientific, Italy) in a dilution ratio of 1:1000 in PBS for 15 minutes and with 1 μg/ml Hoechst 33342 (ThermoFisher Scientific, Italy) for 10 minutes at room temperature. Finally, cells were incubated for 45 minutes with 1 μg/ml of FITC-phalloidin (Sigma Aldrich, Italy), to stain actin cytoskeleton. The images were taken on a A1R+ Nikon (Nikon, Japan) confocal microscope equipped with an apochromat, 20X, NA 0.8 dry objective lens. Images were taken by exciting at *λ=* 488 nm for actin filaments stained with FITC (emission collected through a 525/50 nm band pass filter) and at *λ=* 561 nm for nuclei stained with DRAQ5 (emission collected through a 665/81 nm band pass filter).

Wide-field fluorescence images were taken on a Leica SP5 microscope equipped with a 20X dry objective (HC PL Fluotar, NA=0.5).

### Two-photon Excitation Fluorescence Microscopy

Two photon fluorescence microscopy was performed on a custom setup based on a BX51 (Olympus, Japan) direct microscope equipped with either a water immersion long working distance objective (XL Plan N, 25X, NA=1.05, Olympus, Japan) or a dry objective (Plan N, 20X, NA=0.4, Olympus, Japan). The raster scanning images were acquired through a M610 (ISS Inc., Urbana-Champaign, USA) scanning head coupled to a Ti:Sapph laser (MaiTai DeepSee, Newport, USA) and to two analog photomultiplier tubes (HC125-02, Hamamatsu, Japan).

### Optical Characterization of Microlenses

In all the optical characterization the source was a fiber coupled 635 nm laser diode (HLS635 + P1-630A-FC-1 single mode fiber) collimated by a plan 4X, NA=0.10 objective (Olympus, Japan).

### Wavefront Measurements

The wavefront was measured by means of a Shack-Hartman sensor (WFS20-K1/M, Thorlabs, USA) with a 150 μm pitch array of lenses. A planar Gaussian beam overfills the microlens pupil to be tested. Part of the beam is focused by the microlens on the focal plane of a 63X dry objective that collimates it at its exit pupil (**Fig.SI4.1**). A 4f telescope conjugates the electromagnetic field from the objective exit pupil plane, magnifying it on the entrance pupil of the Shack-Hartman sensor. The fraction of the light that is not captured by the microlens is filtered at the intermediate plane of the 4f telescope by a pinhole (**Fig.SI4.1**), to single out only the wavefront of the beam focused by the microlens.

### Beam profile test

The minimum spot size that can be obtained through the microlenses was measured on an infinity conjugated microscope. The tube lens and the camera sensor (UI-1542LE-M, IDS, Obersulm, D) were carefully aligned to accept rays from infinity, so that the microscope objective collects the electromagnetic field in its front focal plane. The images collected by the camera at various positions of the focal plane of the objective give us the shape of the intensity profile generated by the microlenses.

## Acknowledgments

Part of the experimental characterization was performed at PoliFAB, the micro and nanofabrication facility of Politecnico di Milano (www.polifab.polimi.it). The authors would like to thank the PoliFAB staff for the valuable technical support.

This research has received funding from the European Union under the Horizon 2020 research and innovation program (G.A. No. 964481 – IN2SIGHT).

