## Supplementary materials for "Microlenses fabricated by two-photon laser polymerization for cell imaging with non-linear excitation microscopy"

M. Marini[1,+], A. Nardini[3,+] , R. Martínez Vázquez[2,+], C. Conci[3], M. Bouzin[1], M. Collini[1], R. Osellame[2], G. Cerullo[4], B.S. Kariman[4], M. Farsari[5], E. Kabouraki[5], M.T. Raimondi[3,+,*], G. Chirico[1,+,*].

*1 Department of Physics, Università degli Studi di Milano-Bicocca, Piazza della Scienza 3, 20126, Milan, Italy*
*2 Institute for Photonics and Nanotechnologies (IFN), CNR, Piazza L. da Vinci 32, 20133 Milan, Italy*
*3 Department of Chemistry, Materials and Chemical Engineering "Giulio Natta", Politecnico di Milano, Piazza L. da Vinci 32, 20133 Milan, Italy*
*4 Department of Physics, Politecnico di Milano, Piazza L. da Vinci 32, 20133 Milan, Italy*
*5 FORTH/IESL, N. Plastira 100, 70013, Heraklion, Greece*

+: contributed equally to the work
*: corresponding authors.

**Table of content**

1. **Refraction index of tissues.**
2. **Effect of the inhomogeneity of the propagating medium on the spherical aberrations.**
3. **Microlenses coupled to a raster scanning optical microscope.**
4. **Microlens wavefront measurement.**
5. **Field of view of microlenses.**
6. **Profilometer analysis of the shape of the microlenses.**
7. **Intensity distribution sampling along the optical axis.**
8. **Analysis of the noise and spatial resolution on the images through microlenses.**
9. **On-axis holographic analysis of the profile of the microlenses.**
10. **UV irradiation Optimization during Fabrication Process.**

### SI1. Refraction index of tissues.

-----

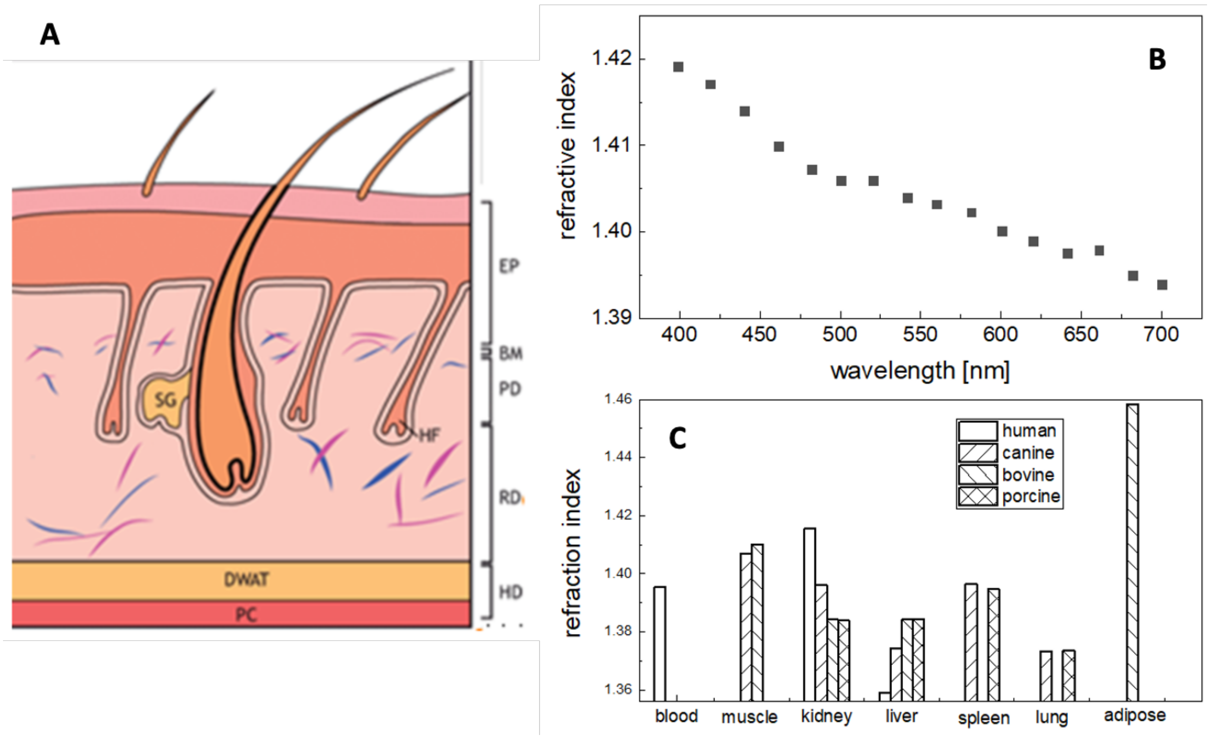


**Figure SI1. A**: The structure of mouse skin. EP: epidermis; BM: basement membrane; PD, papillary dermis; RD, reticular dermis; HD: hypodermis; DWAT: dermal white adipose tissue; PC: panniculus carnosus; HF: hair follicle;SG: sebaceous gland; . Adapted (CC common) from ref. [1]. The skin has a high density of fibroblasts (blue and purple). **B**: Trend of the refractive index of a sample of mouse muscle as a function of the wavelength (open squares), together with the best fits to a linear trend (solid blue line, $y = 1.45 - 7.74 \times 10^{-5}\ \lambda$) and to a power law (solid red line, $y = 1.37 + 180.44\,\lambda^{-1.4}$) (fit on the data taken from ref.[2]). **C**: Refractive index values for various tissues and animals at $\lambda = 633\ nm$ as from ref. [2].

**SI2. Effect of the inhomogeneity of the propagating medium on the spherical aberrations.**

In order to evaluate the effect of the inhomogeneity of the propagating medium on the possibility to tightly focus a collimated laser beam, we have studied an aspheric lens 25 mm in diameter with an effective focal length in air of about 25 mm.
The lens sag surface is given by:

$$z(r) = C * r^2 / (1 + \sqrt{1 - C^2 r^2}) + A_2 r^2 + A_4 r^4 + A_6 r^6$$
$$C = 1/12.6\, mm^{-1};\ A_2 = 3.34 \times 10^{-3}\ mm^{-2};$$
$$A_4 = -2.06 \times 10^{-5}\ mm^{-4};\ A_6 = 5.87 \times 10^{-8}\, mm^{-6}; A_8 = -2.80 \times 10^{-9}\, mm^{-8}$$

The lens is aspherical for propagation in water ($n_1$ =1.34 at $\lambda = 0.8\, \mu m$) for a pupil size of 15 mm. We have then simulated the propagation of light through an inhomogeneous medium composed of a first layer (thickness $d_1$) of water and a second layer (thickness $d_2 = f - d_1$, where $f$ is the lens focal length) of a medium with $n_2 = 1.45$ (**Fig.SI2.1A**). The propagation was computed for a wavelength $\lambda = 0.8\, \mu m$ for a beam coaxial with the optical axis. The phase profiles at the focal plane (minimum confusion circle) for the propagation in pure water and in the inhomogeneous medium ($d_1 = 10\, mm$), for a pupil size of 22 mm, are reported in **Fig.SI2.1B** and **Fig.SI2.1C**, respectively. The aberrations are primarily spherical aberrations and can be described by the Zernike Polynomial with coefficient $a_{4,0}$ . The spherical aberration rises rapidly with the pupil size, as expected[3], and almost doubles its value at 80% of the pupil size when moving from a homogeneous medium to an inhomogeneous medium (see **Fig.1B,** main text). The increase of the spherical aberration coefficient $a_{4,0}$ depends almost linearly on the refraction index changes between the medium layers (**Fig.1B,** main text).
Regarding the minimum spot size of a laser beam in the homogeneous and inhomogeneous cases, we can measure it from the simulated physical propagation of a Gaussian beam 9 mm in waist on the lens. We notice that circular fringes arising from spherical aberrations are evident in the inhomogeneous case (**Fig.SI2.1E**) and prevent a clear measurement of the minimum spot size (beam waist at $1/e^2$ of the maximum intensity). By force fitting the intensity distribution to a Gaussian beam, we can however estimate beam sizes of the order of $5\, \mu m$ and $25\, \mu m$ for the two cases, respectively. This would imply a huge variation of the excitation probability under two-photon regime.

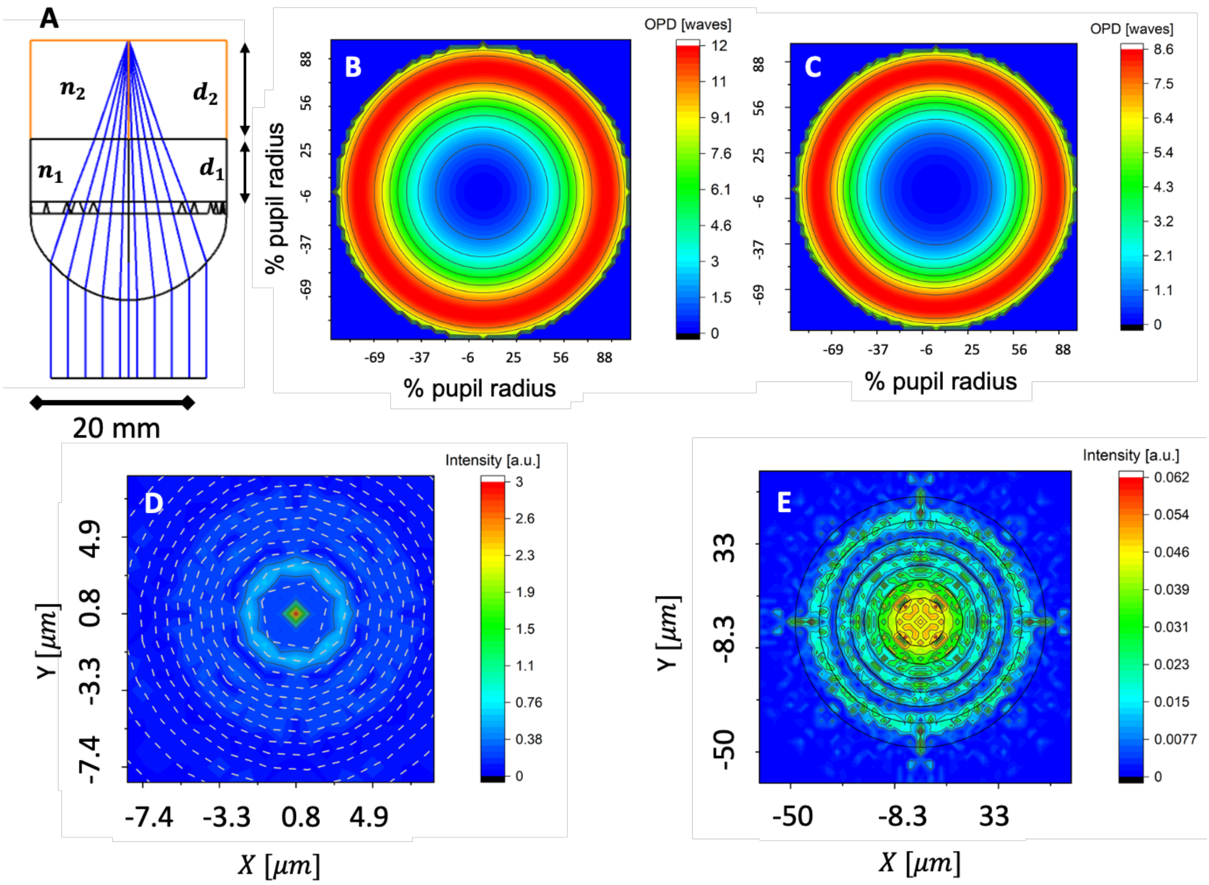


**Figure SI2.1.** Spherical aberrations in inhomogeneous media. **A**: Simulated propagation of a parallel beam focused by a 25-mm focal length, 25-mm diameter, plano-aspherical lens through an inhomogeneous medium composed of a first layer (thickness $d_1$ ) of water and a second layer (thickness $d_2$) of a medium with $n_2 = 1.50$. **B:** Optical path difference (OPD) profiles at the focal plane (minimum confusion circle) for the propagation in the inhomogeneous medium (**B**, $d_1 = 10\,mm$) and in pure water (**C**) for a pupil size of 22 mm. **D-E**: Beam intensity distribution at the focal plane for the homogeneous (**D**) and inhomogeneous ($d_1 = 8\,mm$, **E**) medium. The pupil size was 20 mm.

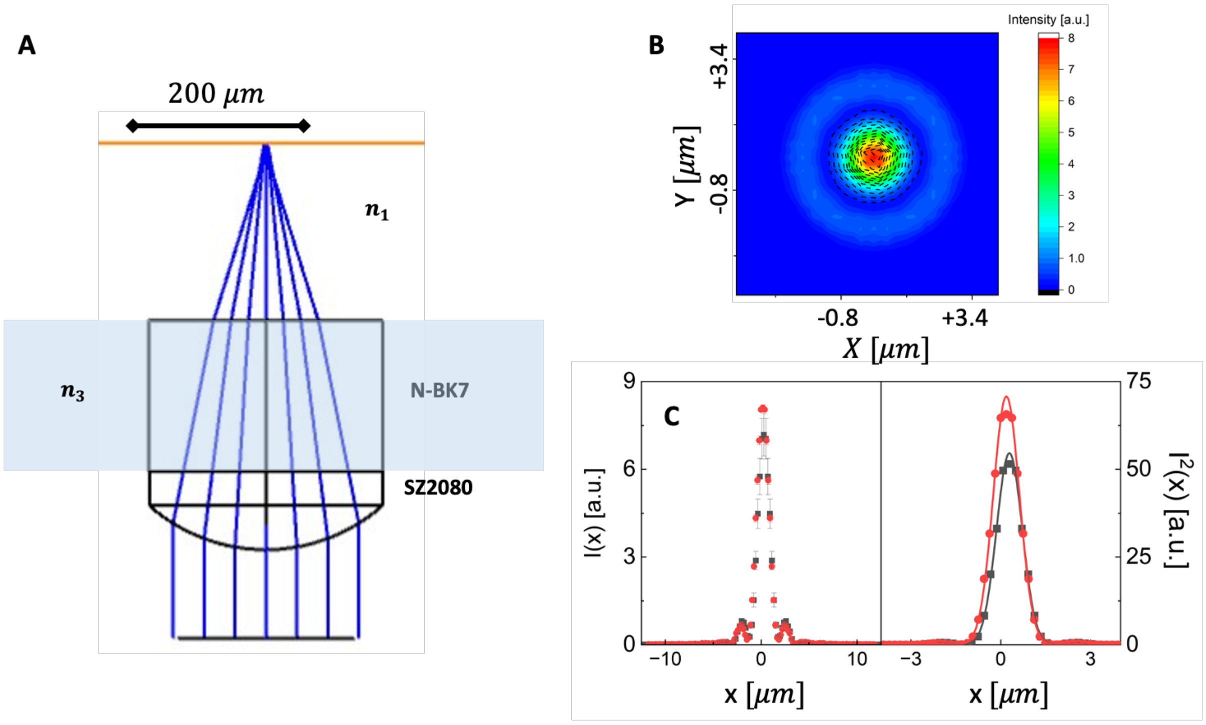


**Figure SI2.2.** Simulation of the beam profile at the focal plane of the microlens (see table I in the main text for the geometrical parameters). **A**: Sketch of the microlens fabricated on a borosilicate glass slide. **B**: Beam intensity distribution at the focal plane of the microlens. **C:** Left plot: profile of the radial intensity of panel B for the case of propagation in air (red symbols) and in water (black symbols). Right plot: profile of the radial dependence of the square of the intensity at the focal plane (panel B) for the case of propagation in air (red symbols) and in water (black symbols) together with the best fit Gaussian distribution (solid lines) whose beam waists are $1.20 \pm 0.03\,\mu m$ and $1.23 \pm 0.03\,\mu m$ for the propagation in air and water, respectively.

**SI3. Microlenses coupled to a raster scanning optical microscope.**

The microlenses can be coupled to an optical microscope in two ways: a finite conjugate setup (**Fig.SI3.1A**) and a virtual image setup (**Fig.SI3.1B**). In both cases we assume to use infinity corrected objectives.

In the first case, the microlens creates a real image of a sample in the focal plane of the objective (**Fig.SI3.1A**). The distances of the sample (p) and of the image (q) planes must be measured with respect to the principal planes and are related to the focal length of the microlens $f_{\mu}$ by $q = pf_{\mu}/(p - f_{\mu})$. The principal planes are at the vertex of the spherical surface and at a fraction of the total thickness (D) of the lens equal to $D/n_L \simeq 258\,\mu m/1.49 \simeq 173\,\mu m$. The thickness is evaluated by summing the thickness of the lens itself and of the borosilicate glass slide and we take an average of the refractive index of the SZ2080 resist ($n_{SZ2080} \simeq 1.46$) and that of borosilicate ($n_{BK7} \simeq 1.52$). The second principal plane lies therefore approximately on the edge between the lens and the glass slide (**Fig. SI3.1A**).

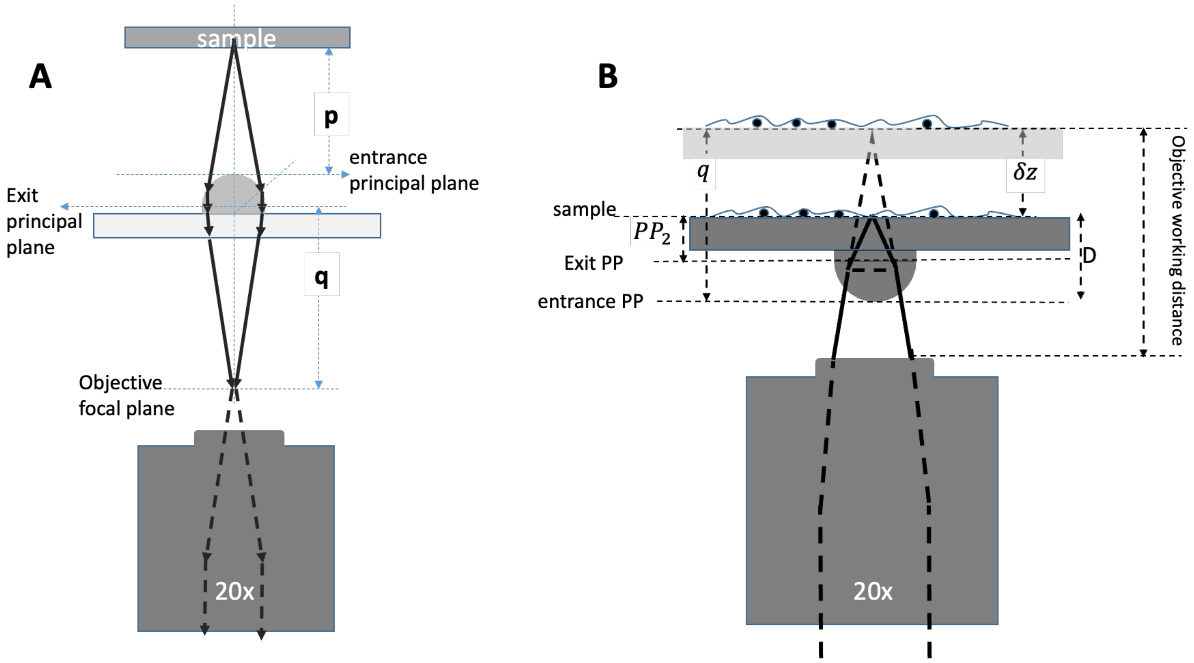


**Figure SI3.1.** Schematic representation of the two possible coupling modes of the microlens with a microscope objective lens. **A**: Finite conjugate coupling between the micro-lens and an objective lens. **B**: The microlens is set at a distance from the exit lens of the microscope objective that is smaller than the objective lens working distance. In this case the microlens creates a virtual image collected by the objective lens. The grayish replica of the sample in the sketch represents the position of the sample for conventional imaging with no microlens in the optical path.

In a second approach the objective lens collects the virtual image of the sample created by the microlens (**Fig.SI3.1B**). In the embodiment realized for this work, the sample (a cell culture) lies on the same glass slide where the microlenses are fabricated. Since the cell culture is approximately bidimensional, the distance p of the sample to the corresponding principal plane is approximately the position, $PP_2$ , of the principal plane with respect to the surface of the coverglass. The distance q of the image plane to the corresponding principal plane corresponds to the full thickness of the microlens device (D in **Fig.SI3.1B**) plus the displacement $\delta z$ by which we need to move the microscope objective to get an image through the microlens. The magnification of the resulting image corresponds to $M = M_T M_{obj}$, where $M_{obj}$ is the objective magnification and $M_T = q/p = (D + \delta z)/PP_2$ is the transverse magnification of the microlens ($PP_2 \simeq 173\ \mu m$).

A simulation of the raster scanning through the microlens compared to the raster scanning without the microlens is reported in **Fig.SI3.2**.

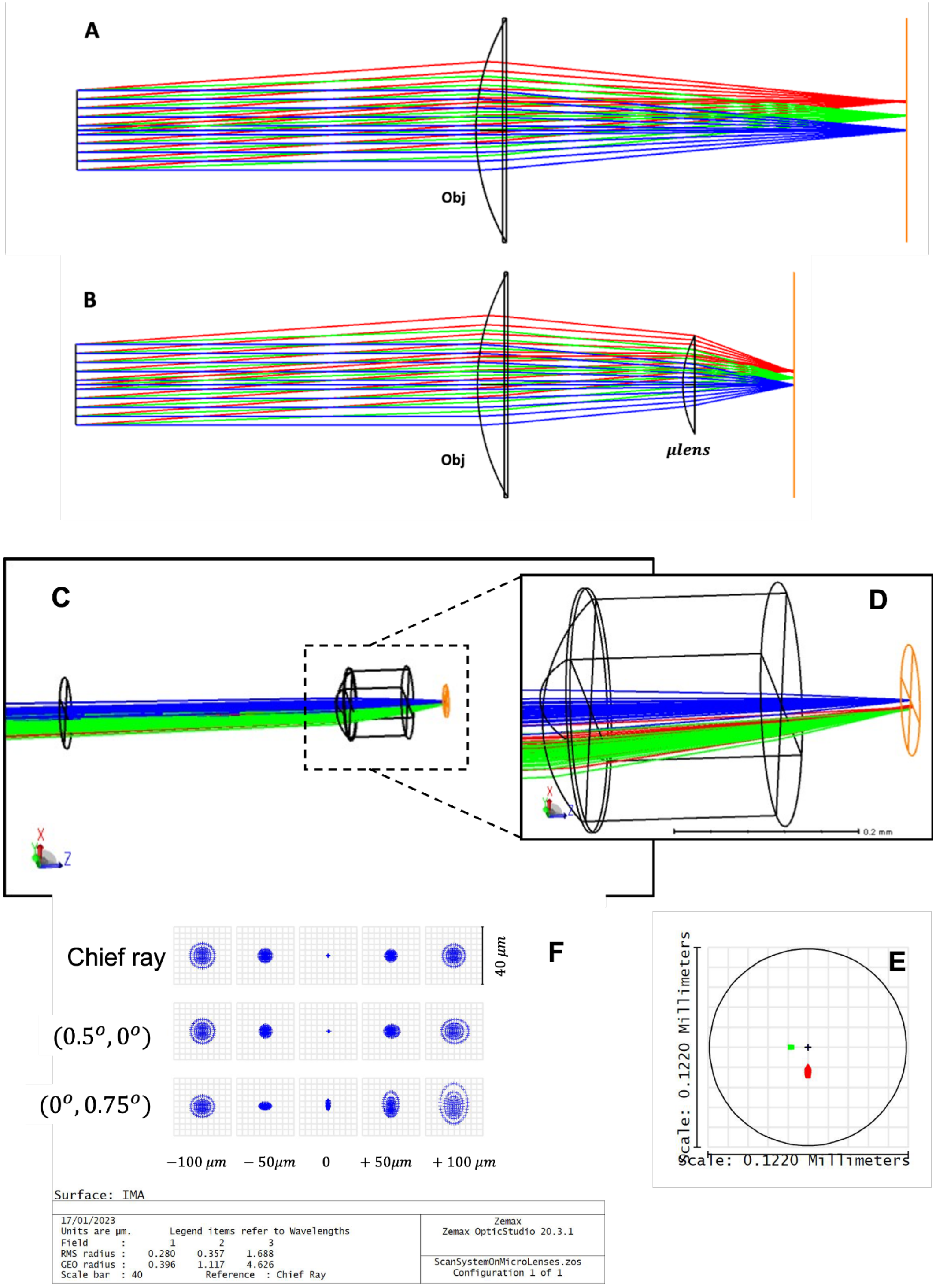


**Figure SI3.2** Raster scanning through a microlens. **A**: Simulation of the raster scanning of a conventional scanning microscope, where the objective lens has been approximated with a single telecentric lens. **B**: Same raster scanning of panel A when a microlens is placed between the objective plane and the sample plane. Note that the microlens increases the overall numerical aperture of the system and consequently reduces the field of view by a factor $M_T$ (calculated above and corresponding to the magnification of the microlens). **C-F**: simulation of the off-axis optical behavior of the microlens ($\lambda = 800nm$). **C:** sketch of the microlens coupled to the glass slide ($170\ \mu m$ thickness) with three fields (chief rays in blue, green and red rays corresponds to $(\alpha_x, \alpha_y) = (0.5^o, 0^o)$ and $(\alpha_x, \alpha_y) =$

$(0, 0.75^{\circ})$, respectively. **D:** blow-up of the dashed box in panel C. **E**: fingerprint plot of the spots on the image plane of the microlens (same color code for the fields). **F**: plot of the spot diagram for the chief ray and the slanted rays through the focal plane in the range $-100\mu m < z < +100\ \mu m$.

We have also characterized the off-axis optical properties of the microlenses by means of simulations. In this case we need to take into account that the microlenses are coupled to a microscope objective, which is a sample space telecentric lens (**Fig.SI3.2A,B**). As reported in **Fig.SI3.2C-F**, over a field of view (on the sample) of about $64\ \mu m$, we find that the RMS spot radius changes from $\simeq 280\ nm$ (on axis) to $\simeq 360\ nm$, at a field of view of $50\ \mu m$. It then increases rapidly to $\simeq 1.6\ \mu m$ at the field of view edge of $64\ \mu m$.

Finally, we consider the case of the use of the microlenses mounted directly in contact with a thick specimen, for example a tissue biopsy, in **Fig.SI3.3**. In this case, by changing the relative position, $z_{O\Pi}$, of the microscope objective lens (exit lens) and the microlens (first principal plane, $PP_1$), we can change at will the position of the object plane in the thick specimen in the range from a few micrometers to a fraction of the focal length $f_\mu$. Taking as a reference the sketch reported in **Fig.SI3.3**, we can derive the effective position of the sample plane imaged by the microlens according to the following relations.

$$p = |q|\, f_\mu/(|q| + f_\mu)) \quad ; \quad p_{eff} = p - T$$

$$q = -(W_d - z_{O\Pi}) \quad ; \quad tan(\alpha) = \Phi/2p \qquad \textbf{(SI3.1)}$$

In these relations the parameters $f_\mu \simeq 240\ \mu m, T \simeq 170\ \mu m, t \simeq 88\mu m, \Phi \simeq 264\ \mu m$ are the microlens focal length, the glass slide thickness, the thickness of the fabricated microlens and the size of the microlens, respectively. $W_d$ is the working distance of the objective. In the following simulations we assume $W_d = 2000\ \mu m.$ From these relations we can compute the position of the focal plane with respect to the glass slide, $p_{eff} = p - T$, and the magnification and the effective $NA = n\, sin(\alpha)$ as a function of the microlens-objective distance $z_{O\Pi}$. As can be seen from **Fig.S3.3B**, the position $p_{eff}$ changes from $44\mu m$ to $\simeq 8\ \mu m$ when $z_{O\Pi}$ changes from $z_{O\Pi} = 2\ \mu m$ to $z_{O\Pi} \simeq 1400\ \mu m$. Notably the maximum extension in depth of the focal plane is limited to $z_{max} = f_\mu W_d/(f_\mu + W_d)$. Therefore we can increase the extension of the imaged volume by increasing the focal length or the working distance of the objective.

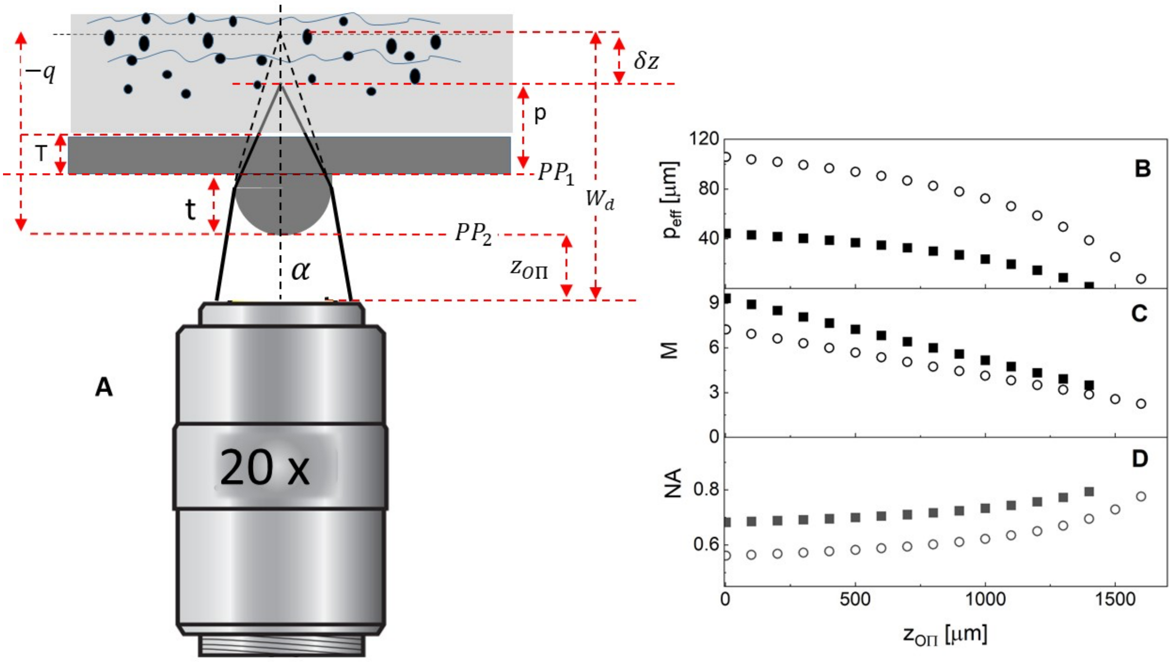


**Figure SI3.3 A**: Scheme of the use of the microlenses coupled to the objective lens to a thick specimen. The relationship between the position ($p_{eff}$) of the effective plane in the sample imaged by the microscope and the relative position of the microlenses and the objective ($z_{O\Pi}$), is given by equations **SI3.1**. The parameters $f_\mu, T, W_d, T, t$ are the microlens focal length, the glass slide thickness, the objective working distance and the thickness of the fabricated microlens, respectively. **B-D**: trend of the effective focal plane position $p_{eff} = p - T$, the absolute value of the magnification M and the numerical aperture of the microlens $NA = n\,sin(\alpha)$, where n = 1.33, as a function of the microlens-objective distance $z_{O\Pi}$. Filled and open symbols in each panel correspond to the case $f_\mu = 240\ \mu m$ (filled symbols) and $f_\mu = 340\ \mu m$ (open symbols).

**SI4. Microlens wavefront measurement.**

The wavefront was measured by means of a Shack-Hartman sensor (WFS20-K1/M, Thorlabs, USA) with a 150 μm pitch array of lenses. The optical configuration adopted (**Fig.SI4.1**) allows us to test the lenses by focusing a collimated beam. A planar Gaussian beam impinges on the microlens to be tested with a $1/e^2$ size of about 9 mm. Part of the beam is focused by the microlens on the focal plane of a 63X dry objective (Zeiss A-Plan 63x, NA= 0.8). The electromagnetic field is sampled by the 63X objective lens and collimated at its exit pupil. A 4f telescope conjugates the electromagnetic field from the objective exit pupil plane, magnifying it on the entrance pupil of the Shack-Hartman sensor. The fraction of the light that is not captured by the microlens is filtered at the intermediate plane of the 4f telescope by a pinhole (PH in **Fig.SI4.1**). In this way it is possible to single out only the wavefront of the beam focused by the microlens.

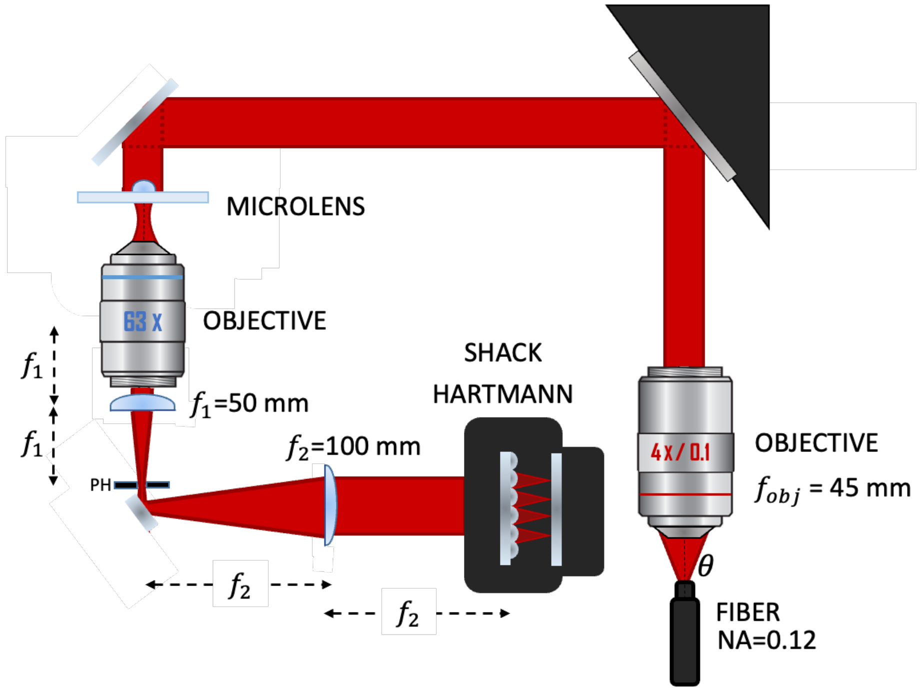


**Figure SI4.1**. Sketch of the optical setup adopted for the wavefront measurement.

Comparison to theoretical treatments. The microlenses were always tested in the configuration shown in **Fig.SI4.1**, this condition providing minimal spherical aberration[3] and corresponding to a shape factor $Q = (R' + R)/(R' - R) = +1$, where R' and R are the curvature radii of the two lens surfaces, and an imaging factor $P = 1 - 2q/f$, where $f$ is the lens focal length (ref.[3], p.243]. The prediction of the spherical aberration *of a thin lens* with the Seidel description of the wavefront[3] is $C_{40} = g(n_L)\, a^4/f^3$, where $a$ is the pupil radius and $g(n_L, P, Q)$ is a correction factor that depends on the refraction index of the lens (for $n_L \simeq 1.46$, P=-1, Q=+1, $g \simeq 0.34$). We notice that this expression is valid for a thin lens and that it is based on a description of the wavefront in terms of powers of the pupil radial coordinate, and not in terms of the Zernike polynomials.

The Shack-Hartman (SH) sensor and Zemax software adopt instead a description of the wavefront in terms of the Zernike polynomials, that are labeled according to the Noll's convection[4] and normalized to their corresponding root mean square (RMS) values. In this way, the contribution of each polynomial to the wavefront RMS is 1 (for the spherical aberration polynomial, ${Z_4}^0$, the RMS value is $\sqrt{5}$). Therefore, the spherical aberration amplitude (i.e. the coefficient of ${Z_4}^0$ in the description of the wavefront) measured by the SH sensor, ${a_{4,0}}^{(SH)}$ should be equal to the one provided by Zemax, ${a_{4,0}}^{(Z)} = {a_{4,0}}^{(SH)}$. If we wanted to compare the amplitude of the spherical aberration predicted by the Seidel description (see ref.[3], Eq. 4.88), ${a_{4,0}}^{(S)}$, to the one given in terms of the Zernike polynomials, ${a_{4,0}}^{(Z)}$, we would need to take into account an additional normalization factor $6/\sqrt{2}$ (see ref. [5] Eq.9 ch. 9.3): ${a_{4,0}}^{(S)} = 6/\sqrt{2}\, {a_{4,0}}^{(SH)}$.

For a *microlens* with the parameters reported in **Table I** (main text), the Seidel prediction is ${a_{4,0}}^{(S)} \simeq 2.8\, \mu m$ that, converted in Noll's notation, is equivalent to an amplitude $\simeq {a_{4,0}}^{(S)}/4.25 \simeq 0.66\, \mu m$. For comparison, the measured value for the microlens is ${a_{4,0}}^{(SH)} = 0.10 \pm 0.01\, \mu m$ (averaged over 3 different microlenses) and the Zemax prediction is ${a_{4,0}}^{(Z)} = 0.15\, \mu m$. This is 50% higher than the measured value, but both values are 4-5 times larger than the Seidel prediction, which is an approximation for a thin lens and it is not sufficient to recapitulate the observations. Indeed, in the experimental evaluation of the aberrations, more than the amplitude of the individual Zernike components describing the wavefront, it is the wavefront RMS value that can be compared to theoretical predictions. While theoretically the distribution of the Zernike weights is typically well peaked on a specific aberration component (like here on the spherical aberration), experimentally the wavefront is described by a distribution of Zernike polynomials and among them the spherical component, $a_{4,0}$, is the dominant one. As a confirmation of this, we observe that the measured average wavefront RMS is $0.13 \pm 0.02\ \mu m$ that is in good agreement with the Zemax prediction, RMS $= 0.15\ \mu m$. As a comparison to a conventional plano-convex lens, we employed a Thorlabs LA1252 lens with front focal length f=25 mm and 25.4 mm diameter. By using a pupil of about 9 mm size at the entrance of the lens we measured $a_{4,0}{}^{(SH)} \simeq 0.17 \pm 0.02\ \mu m$ and an RMS value $\simeq 0.17\ \mu m$, that compares very well with the RMS value of 0.18 $\mu m$ of the simulated wavefront. However, again, the comparison of the amplitude of the spherical mode computed with the Seidel prediction (see ref.[3], Eq. 4.88] disagrees by a factor of 4 with the measured value, since $a_{4,0}{}^{(S)} \simeq 4.4\ \mu m$ and therefore $6/\sqrt{2}\ a_{4,0}{}^{(SH)} \simeq 0.7\mu m\ \simeq 4a_{4,0}{}^{(SH)}$ .

We finally tested the Shack-Hartman setup by measuring the change of the spherical aberration amplitude $a_{4,0}$ as a function of the pupil size for the macroscopic plano-convex lens finding, to a good degree of accuracy, the expected fourth power increase with the pupil size, as demonstrated in **Fig.SI4.2** (solid line).

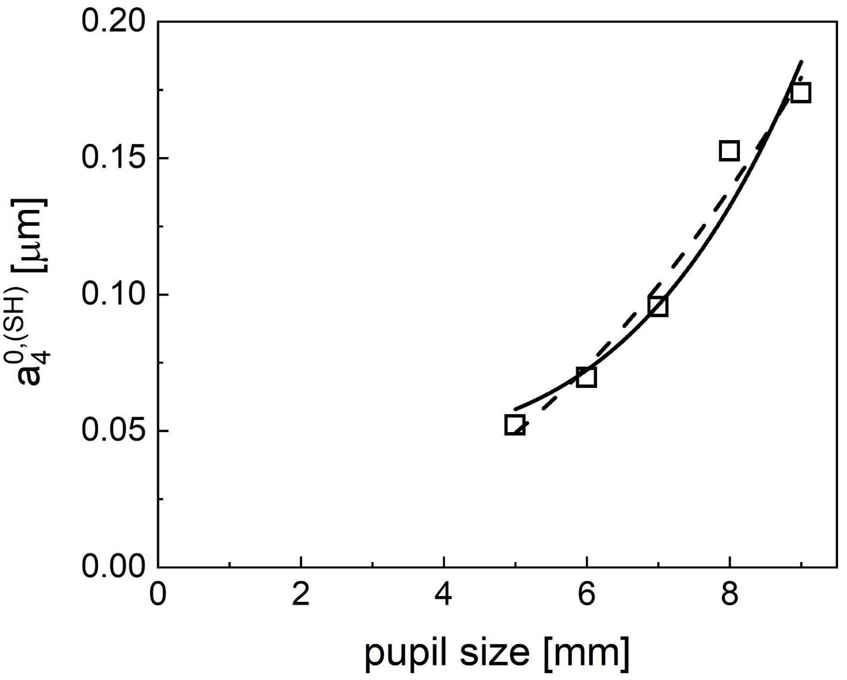


**Figure SI4.2** Measurement of the spherical aberration coefficient for a plano-convex lens (Thorlabs, LA1252) as a function of the pupil size. The solid and the dashed lines are the best fits of the trial function $a_{4,0}(p) = A + Bp^m$ to the data with best fit parameters $[A, B, m] = [0.044\,\mu m; 2.1 \times 10^{+6}\,\mu m^{-3}, 4]$ (solid line) and $[A, B, m] = [0\,\mu m; 3.1 \times 10^{+3}\,\mu m^{-1.2}, 2.2]$ (dashed line).

### SI5. Field of view of microlenses

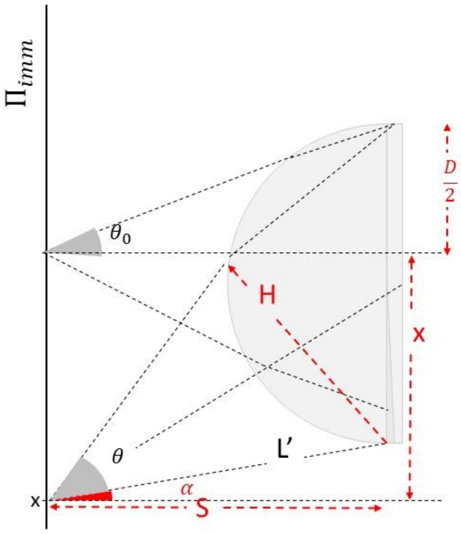


$$\tan(\theta_0) = \frac{D}{2S}$$

$$\tan(\alpha) = \frac{x - \frac{D}{2}}{S}$$

$$\tan(\alpha + \theta) = \frac{x + \frac{D}{2}}{S}$$

$$L' = \sqrt{\left(x - \frac{D}{2}\right)^2 + S^2}$$

$$H = L' sen(\theta)$$

$$P(\theta) = P_0 \left(\frac{D}{H}\right)^2$$

**Figure SI5.1** Computation of the field of view of a microlens under wide field imaging. The power collected as a function of the angular position on the sample plane $\Pi_{sample}$ is computed as $P(\theta) = P_0\,(D/H)^2$.

The field of view of a microlens under wide field imaging is computed according to the scheme reported in **Fig. SI5.1**. The power collected as a function of the angular position on the image plane is computed as $P(\theta) = P_0\,(D/H)^2$. The field of view is defined as the angular position on the sample plane $\Pi_{sample}$ where the collected power is 37 % of the maximum collected power on the optical axis. A simulation for different values of the distance S is reported in **Fig.SI5.2**.

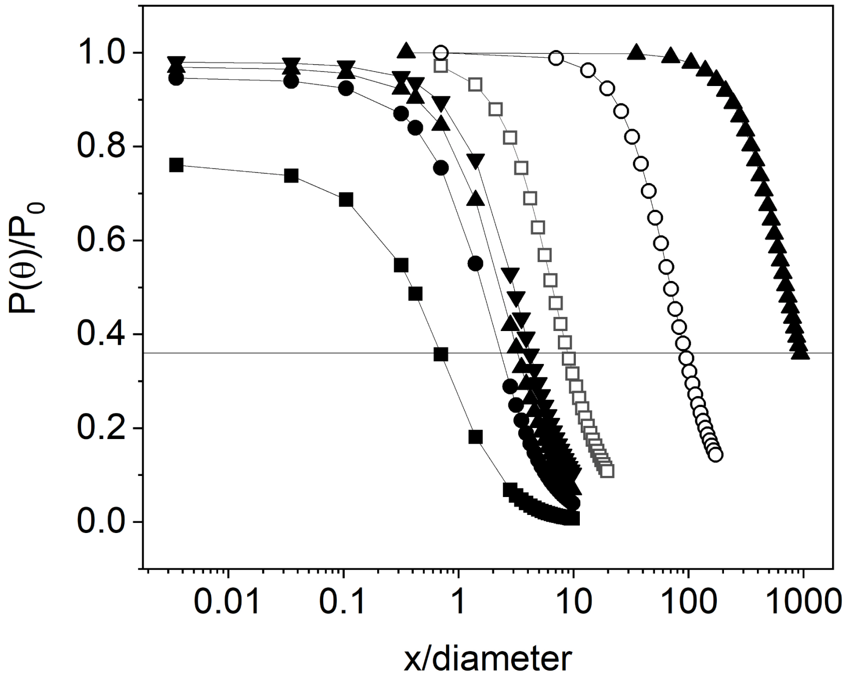


**Figure SI5.2.** Change of the fraction of the collected power as a function of the angular position on the sample plane for the plano-convex microlens with front focal length $f_{FFL} = 164\ \mu m$. The solid horizontal line indicates the 37% of the maximum power. The symbols correspond to different values of the sample-lens distance $S$. The lens is approximated as a thin lens for this evaluation: $S = 260\mu m$ (filled squares), $600\ \mu m$ (filled circles), $800\ \mu m$ (filled up-triangles), $1\ mm$ (filled down-triangles), 2 mm (open squares), 2 cm (open circles), 20 cm (filled up-triangles).

**SI6. Profilometer analysis of the shape of the microlenses.**

The overall shape of the fabricated microlenses was analyzed by using a surface profilometer (KLA Tencor P-17) which is based on a stylus-based scanning, using a 2.5 µm stylus tip radius. Excluding from the analysis the set of points affected by the footprint of the stylus (from the ground level, about 45 µm along the vertical axis of the lens), the surface profile of the 2PP microlenses fitted the spherical function with a R-square of 0.998 (**Fig.SI6**). The average height and curvature slightly differ from the model (see **Table I** in the main text) and they respectively are 93 µm (+5%) and 193 µm (-3.5%, **Fig.SI6B**). A deviation of the fabricated surface from the desired spherical one is expected due to the dimensions of the polymerized voxel and some possible shrinking occurring upon developing the microstructure. This deviation can be corrected by working on the CAD shape of the lens and the stability can be improved by increasing the thickness of the 2PP shell by multi-scan, specially for the pedestal which is the weakest part of the lens.

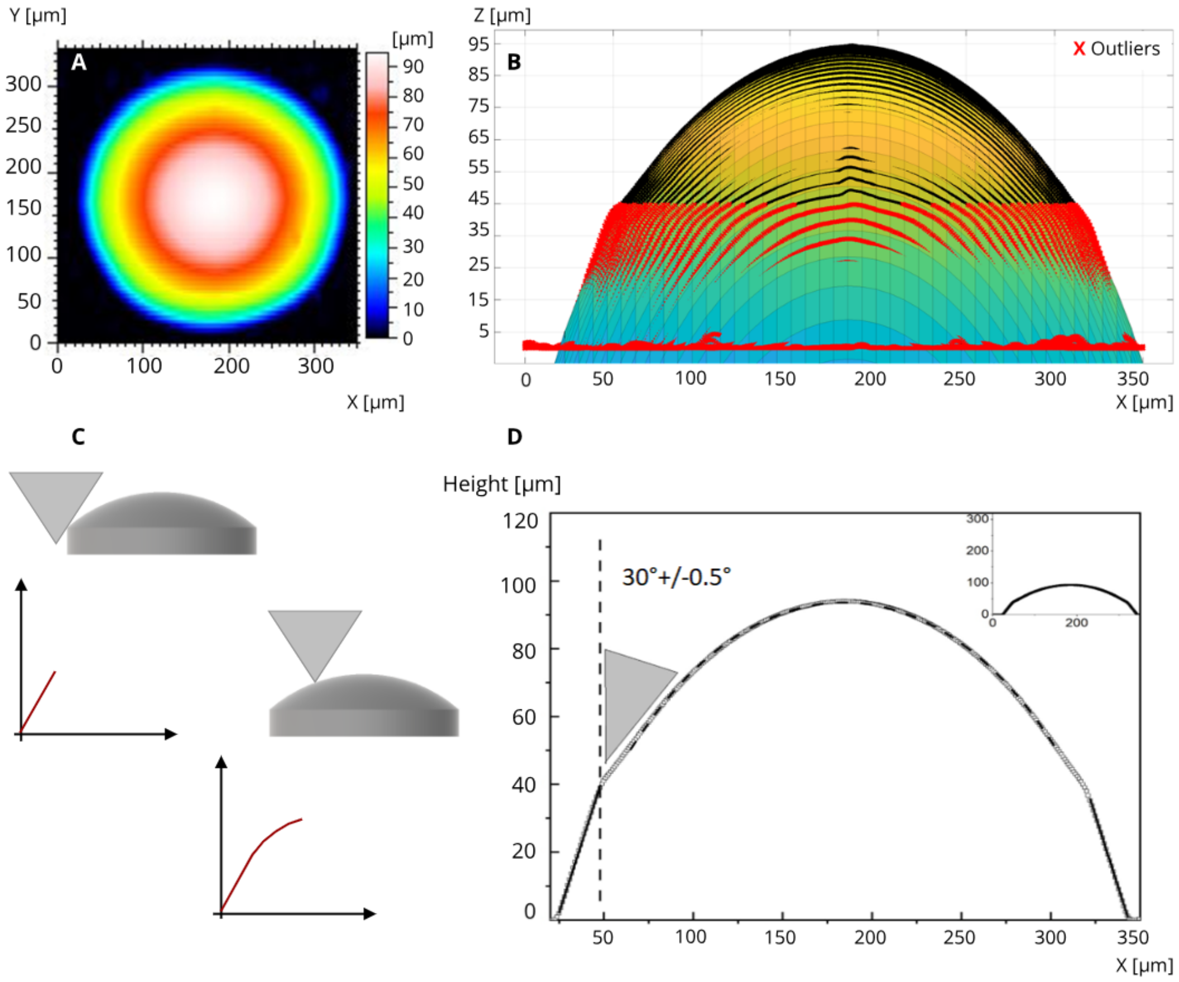


**Figure SI6.** Outcome from the profilometry measurements. **A**: Height of the overall lens plotted using a color code diagram; **B**: Spherical fitting of the surface profile of the fabricated lenses excluding the data affected by the footprint of the stylus tip of the profilometer (red points). **C**:

sketch of the convolution of the actual surface by the tip of the profilometer. **D:** Spherical fitting of the surface profile of the fabricated lenses excluding the data affected by the footprint of the stylus tip of the profilometer, that are instead fit to two linear trends that correspond to a slope with an angle of $30^o \pm 0.5^o$, compatible with the nominal full tip angle $\simeq 60^o$. The best fit of the curvature radius is $R = 193 \pm 4\, \mu m$.

### SI7. Intensity distribution sampling along the optical axis.

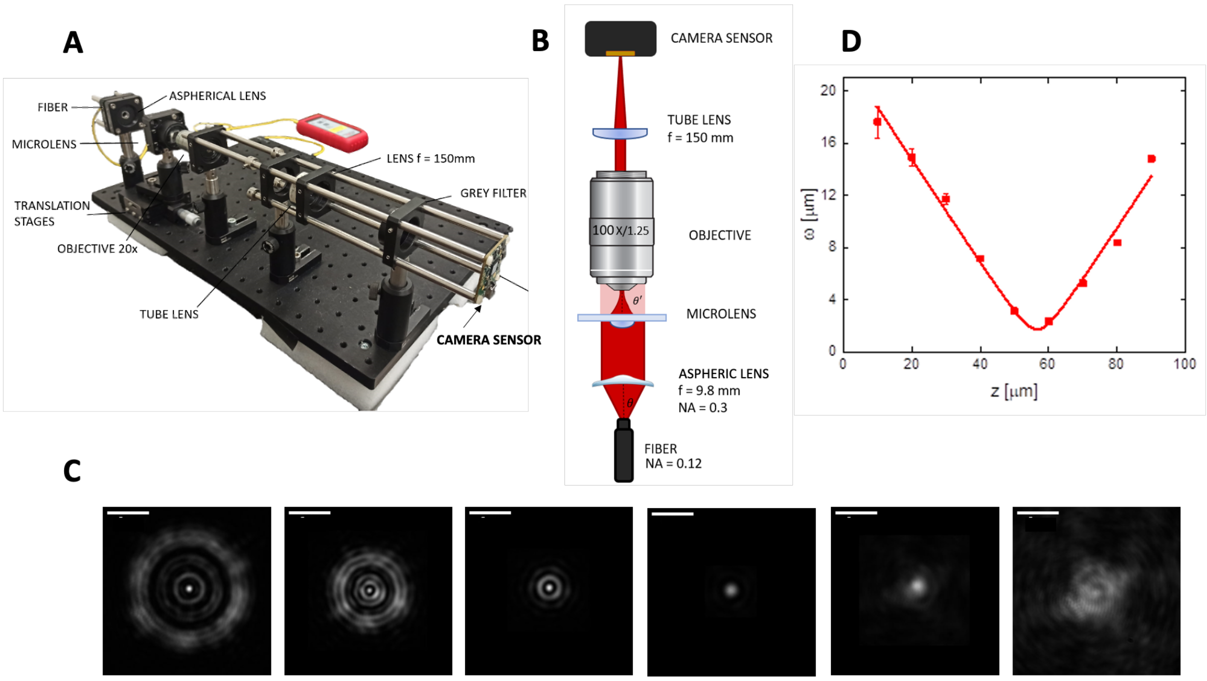


**Figure SI7.** Minimum Spot Size test. **A**: Photograph of the setup for the test of the minimum beam spot and the Rayleigh range of the microlenses. **B**: Sketch of the optical path. The tube lens and the camera sensor are aligned to accept rays from infinite. **C**: Images of the intensity profiles collected at various distances along the microscope objective optical axis. The mutual distance between the planes is 10 μm, starting from the plane closer to the microlens (from the left to the right). Scale bars 5 μm. **D**: Trend of the effective beam waist measured by the cumulative sum of the intensity profiles in panel **C** fitted to **Eq.SI7.3**. The solid line corresponds to the fit of the effective beam waist data to **Eq.SI7.2**. The best fit values are $w_0 = (2.0 \pm 0.3)\ \mu m$ and $z_R = (5.6 \pm 1.5)\ \mu m$ (average values over a full array of 7 microlenses).

The intensity distribution on planes perpendicular to the optical axis provides a direct visualization of the effect of the spherical aberrations[5]. We have sampled the intensity at $10\ \mu m$ steps by means of an infinite conjugated microscope working at 635 nm (**Fig. SI7**). The tube lens and the camera sensor are aligned to accept rays from infinity. In this way the microscope objective collects the electromagnetic field in its front focal plane (**Fig. SI7A-B**). The images collected by the camera at various positions of the objective focal plane give us the shape of the intensity profile generated by the microlenses. An exemplary set of images is reported in **Fig.SI7C**, evidencing the effect of the spherical aberrations of the microlens.

Due to the presence of the spherical aberrations, a measurement of the value of the $1/e^2$ beam size cannot be directly obtained by fitting the intensity distributions of **Fig.SI7C**. At each plane perpendicular to the optical axis (for each image in **Fig.SI7C**), we can try to obtain an approximate value of the $1/e^2$ beam size by computing and fitting the cumulative sum of the beam intensity distribution along the columns of the image. In fact, an ideal Gaussian beam is determined by the beam waist, $w_0$, and the Rayleigh range, $z_R = \pi w^2{}_0/\lambda$, as:

$$I(x,y,z) \; = A/w(z)^2 \, exp\{-2[(x-x_0)^2 - (y-y_0)^2\,]/w(z)^2\,\} + B \qquad \text{(SI7.1)}$$

$$w(z) = w_0 \sqrt{1 + (z/z_R)^2} \qquad \text{(SI7.2)}$$

Given the image of the beam intensity profile, the cumulative sum of the intensity distribution detected along the image columns (here the y axis) is given by ref.[6]:

$$< I(x,z) >_y = \; A\pi w^2(z)[1 - erf(\sqrt{2}(x_0 - x)/w(z)] \; + \; BMx \qquad \text{(SI7.3)}$$

Here $M$ is the length in pixels of the image columns. By the computation of $< I(x,z) >_y$ for each image in **Fig.SI7C** and its fit to **Eq. SI7.3** we obtain an effective value of the $1/e^2$ beam radius $w(z)$ at each value of the distance, z, along the optical axis. $w(z)$ is plotted as a function of z in **Fig.SI7D**. The best fit values of the beam waist and the Rayleigh range obtained by fitting these data to **Eq. SI7.2** are $w_0 = \; (2.0 \pm 0.3)\,\mu m$ and $z_R \; = (5.6 \pm 1.5) \; \mu m$. As a comparison, the value of the Rayleigh range that can be predicted from the beam waist measured from the FWHM on the minimum spot size plane (**Fig.4D** in the main text) is $z_R \; \simeq (2.5 \pm 1.1) \; \mu m$. The approximate value $z_R \; = (5.6 \pm 1.5) \; \mu m$ appears therefore overestimated, confirming that the presence of side lobes due to the spherical aberrations in the imaged spot profiles prevent exploiting the Gaussian modeling of the beam. Even the analysis based on the cumulative sum of **Eq. SI7.3** fails in providing an accurate estimate of $z_R$, that should be measured by the FWHM on the minimum spot size plane as performed in the main text.

### SI8. Analysis of the noise and spatial resolution on the images through microlenses.

Quality assessment on the images. In order to evaluate the quality of the images acquired through the microlenses with respect to the ones taken through the microscope objective only, we can exploit two algorithms, one based on a perception based metrics and the second based on the computation of the standard deviation of the digital level in the images. In the first case we exploit "Perception based Image Quality Evaluator" (PIQE).[7] This algorithm, which belongs to the class of *No-Reference Image Quality Assessment* algorithms, provides a score in the interval [0-100], where the best quality image is assigned a PIQE = 0. The application of the PIQE algorithm to the images reported in **Fig.6** is summarized in table SI8.1. for the images reported in Fig.6 of the main text.

| **Table SI8.1: evaluation of signal/noise metrics on the confocal and non-linear excitation images.** | | |
|---|---|---|
| **Image** | **PIQE score** | $\frac{\sigma_F}{< F >}$ |
| confocal image, control (Fig.6D) | 30.5 | $0.17 \pm 0.03$ |
| confocal image, through microlenses (Fig.6C) | 50.0 | $0.22 \pm 0.06$ |
| TPE image, control dry objective 20X (Fig.6G) | 62.4 | $0.32 \pm 0.05$ |
| TPE image, control water immersion objective 25X (Fig.6I) | 61.1 | $0.22 \pm 0.01$ |
| TPE image through $\mu$lenses, dry objective 20X (Fig.6F) | 61.0 | $0.26 \pm 0.08$ |
| TPE image through $\mu$lenses, water objective 25X (Fig.6H) | 61.6 | $0.20 \pm 0.03$ |

The quality of the images can also be evaluated directly on their digital content by means of the standard deviation of the distribution of digital levels, $\sigma_F$. We selected ROIs of identical size (22x22 pixels) on regions of uniform signal and computed the ratio of the standard deviation $\sigma_F$ ,divided by the average value of the fluorescence signal, $\frac{\sigma_F}{<F>}$ . The result of this analysis is summarized in table SI8.1 and in **Fig. SI8.1**.

The *48%* change in the perceptive PIQE score between the confocal reference image and the one collected on the confocal microscope coupled to the microlenses, indicates that in the confocal case the degradation of the figure quality when using the microlenses is substantial. This is likely due to a reduction of the total collected signal due to a misalignment with respect to the confocal pinhole. On the other hand, for the case of non-linear excitation the PIQE score does not change substantially among the various configurations tested. However, the relative standard deviation analysis on uniform ROIs taken from **Fig.6F-I** (corresponding to two-photon excitation) indicates instead that the images collected through the microlenses under non-linear excitation have similar or even better metrics (lower value of $\frac{\sigma_F}{<F>}$) with respect to their corresponding control cases. The result of this analysis is reported in **table SI8.1** (average values) and in **Fig. SI8.1**. Both in the case of the 20x dry objective and of the 25x water immersion objective, the value of $\frac{\sigma_F}{<F>}$ is similar for the control and the microlens acquisition.

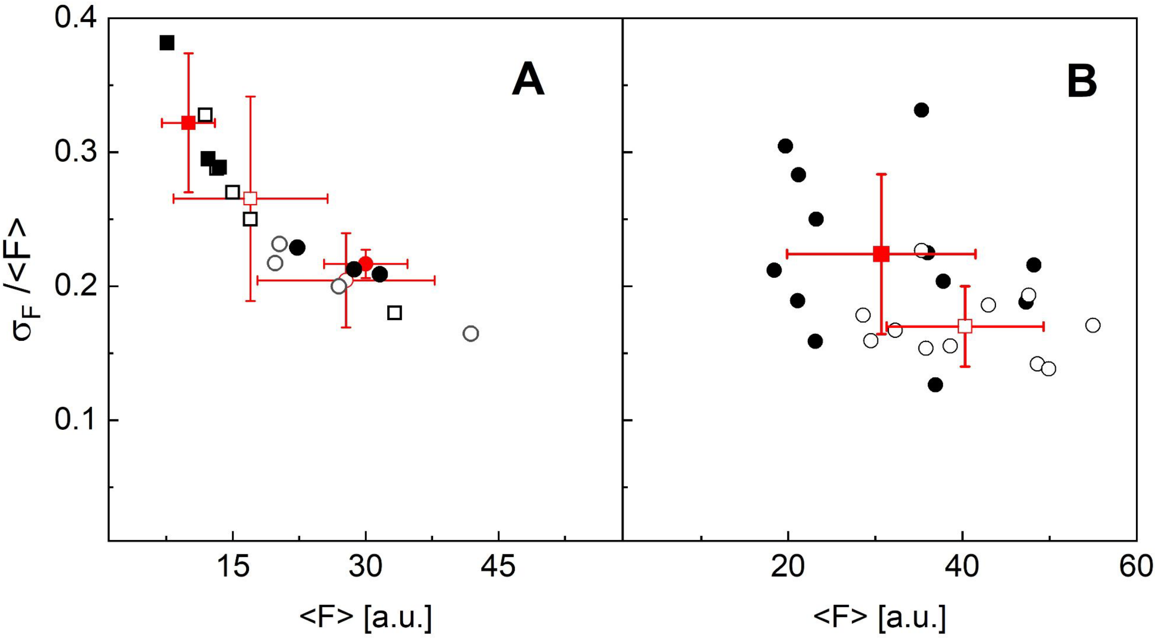


**Figure SI8.1**. Result of the analysis of the relative RMSE (root mean square error) on the distribution of color levels in 22x22 pixels size ROI in the images reported in Fig.6. **Panel A**, analysis of the TPE images in **Fig.6**: **Fig.6F**, corresponding to the case of the water immersion objective (25X) coupled to the microlenses (open black circles and open red circle); **Fig.6G**, corresponding to the case of the water immersion objective (25X) only (filled circles and filled red circle); **Fig.6H**, corresponding to the case of the dry objective (20X) coupled to the microlenses (open black squares and open red square); **Fig.6I**, corresponding to the case of the dry objective (20X) only (filled black squares and filled red square). The red symbols correspond to the average value of the relative RMSE for each subset. **Panel B**, analysis of the confocal images in **Fig.6**: open black circles and open red square correspond to the case of the confocal microscope coupled to the microlenses (**Fig.6B**); filled black circles and filled red square correspond to the case of confocal microscopy with no microlenses coupled to the objective (**Fig.6D**).

<u>Estimate of the effective spatial resolution on the images</u>. We cannot measure the spatial resolution directly on the cell samples. However, we can estimate it by measuring the minimum size of the features on the images. For the confocal images, where the actin filaments can be segmented well, we have drawn spatial profiles and fit them to multi-peaks Gaussian trial functions (**Fig. SI8.2**). We find that that the average size of the features (likely thin bundles of actin filaments) are $\delta x = 0.8 \pm 0.2\ \mu m$ and $\delta x = 1.0 \pm 0.3\ \mu m$ for images taken respectively on the confocal microscope and through microlenses coupled to the confocal microscope.

For the case of TPE images, we exploited the chromatin inhomogeneities in the cell's nuclei to give an estimate of the spatial resolution. Inhomogeneous lumps of signal in the nuclei were fit to 2D Gaussian multi-component functions, finding a radial $1/e^2$ radius $< \delta r >= 2.3 \pm 0.6\ \mu m$ and $< \delta r >= 2.0 \pm 0.6\ \mu m$ for the TPE microscope (water immersion objective) and for the case in which the microlenses were coupled to the same TPE microscope objective (see **Fig. SI8.3** for details on this analysis).

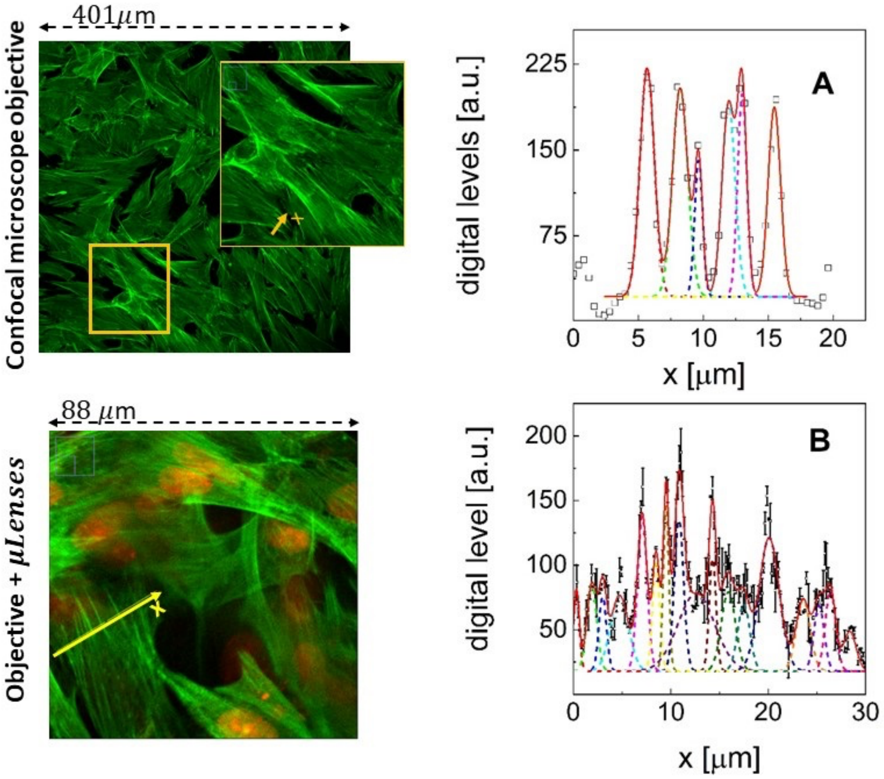


**Figure SI8.2**. Analysis of the spatial resolution on confocal images taken through the microscope objective only (**panel A**), or through the microscope objective coupled to the microlenses (**panel B**). The spatial profile is taken along the x axis sketched on the images (or the inset). The profiles were analyzed by fitting to multipeak Gaussian trial functions as reported in the graphs. The solid lines are the cumulative fit of the profiles. The short-dashed lines are the individual Gaussian components. The average values of the $1/e^2$ half widths ($w_{1/e^2}$) of the Gaussian components ($FWHM \simeq 1.17\ w_{1/e^2}$) are $\delta x = 0.8 \pm 0.2\ \mu m$ and $\delta x = 1.0 \pm 0.3\ \mu m$ for panel A and panel B, respectively.

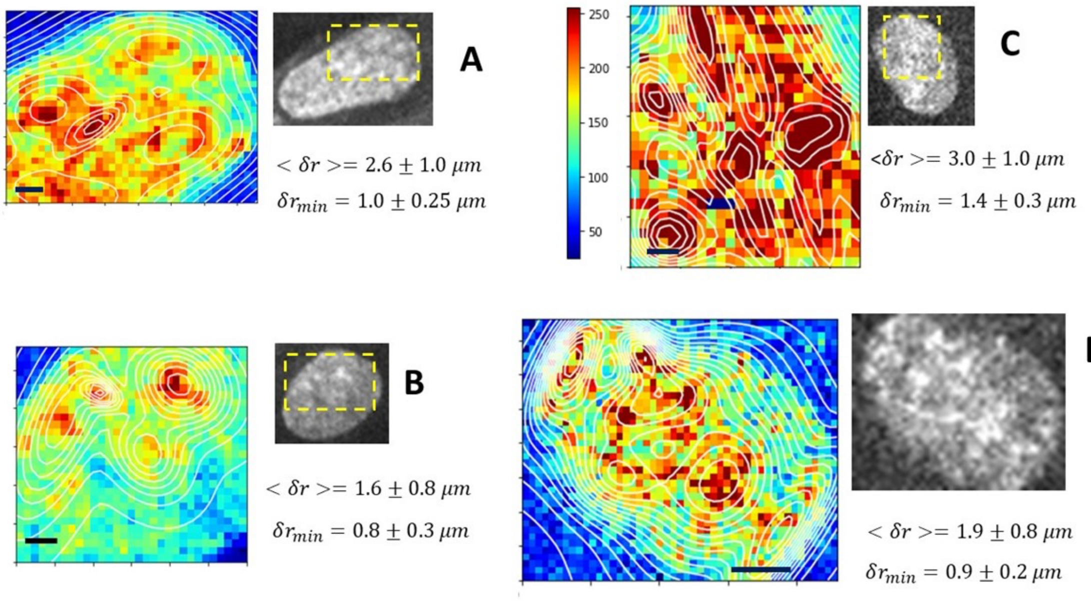


**Figure SI8.3** Analysis of the effective spatial resolution on TPE images for the case in which the microlenses were (**panel A, B**) or were not (**panel C, D**) coupled to the objective of the non-linear excitation microscope. ROIs (size from $7x7\ \mu m^2\ to\ 18x18\ \mu m^2$) were fit to multi-peak 2D Gaussian trial functions. The average and the minimum values of the radial width $\delta r = \sqrt{w_{1/e^2,x}{}^2 + w_{1/e^2,y}{}^2}$ of the 2D Gaussian components are reported in the panels. ROIs from the images (in grayscale) are reported in jet colormap superimposed to the best fit multi-component Gaussian best fit functions (contours in white). The bar in the contour images corresponds to to the size of $1\ \mu m$.

### SI9. On-axis holographic analysis of the profile of the microlenses.

The measurement of the wave front made through a Shack-Hartman (SH) sensor is suitable for the measurement of the aberrations for which not many sampling points are necessary. In our case, the number of sampling points measured by means of the SH sensor is enough to fit the wave front to Zernike polynomials at least up to the 15th order. Moreover, a SH sensor can be easily used as the source of a feedback loop for the wave front aberrations corrections of the microlenses by means of a deformable mirror.

A direct measurement of the profile of the microlens can also be performed by means of digital holography microscopy (DHM). As described recently for the characterization of metalenses, off-axis DHM[8] allows the reconstruction of the whole profile of the lens. Once the full field phase and amplitude is reconstructed, one can compute all the optical response functions.[8] Since, the aberrations of the microlenses are already characterized by means of Zernike analysis of the wave front measured by means of the SH sensor, we limit ourselves here to the implementation of the on-axis DHM, as outlined in **Fig.SI9.1**.

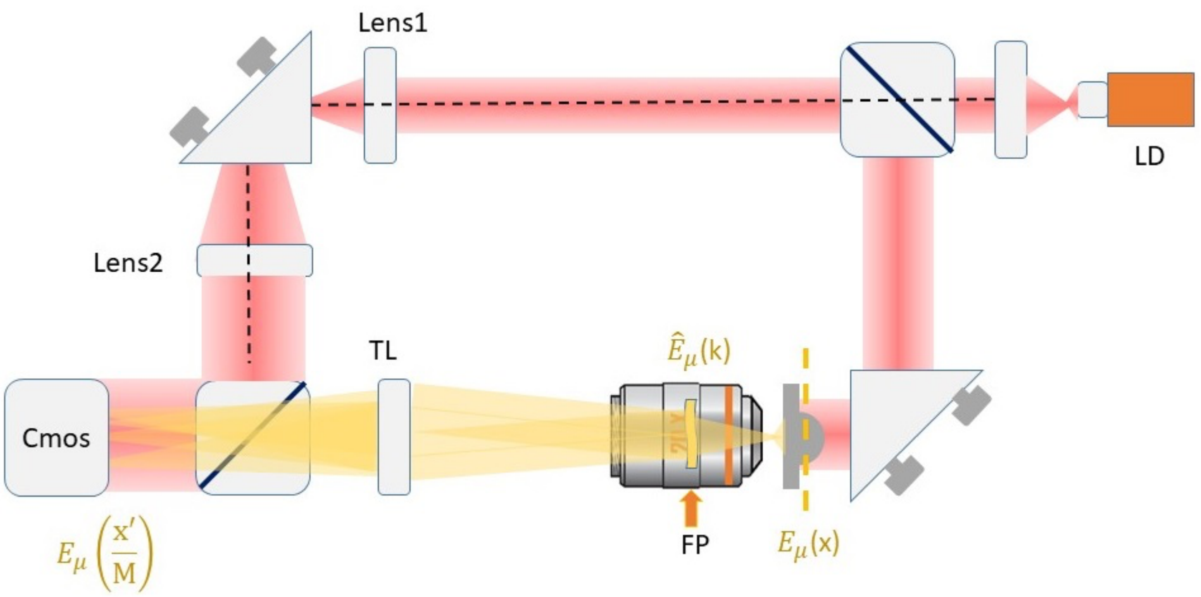


**Figure SI9.1.** Outline of the on-axis DHM setup used for the measurement of the iso-OPD lines on the microlens profile. It is based on a Mach-Zender interferometer. The light from a laser diode ($\lambda = 635\ nm$) is collimated and expanded (lens1-lens2 telescope) in the top path, to fit the size of the beam collected by the tube lens (TL) on the bottom path. The light field from the the dashed plane $E_\mu(\overrightarrow{x})$ is Fourier transformed in the objective Fourier plane (FP) and conjugated again on the CMOS camera (Cmos) superimposed with the reference beam.

This technique allows us to sample the microlens profile at $2\pi$ spacing in the optical path. We can therefore retrieve the positions $x_k$ at which the phase shift $\Delta\Phi(z_k, x_k)$ with respect to the reference wave front is equal to an integer number of $2\pi$: $\Delta\Phi(z_k, x_k) = 2\pi k$ (k=1,2…), and compare them to those expected from the optical design. The image of the lens appears then superimposed on the sets of lines of equal optical path difference (iso-OPD lines, see **Fig.SI9.2A**). The pattern of the fringes can be digitized and averaged over the polar angle as reported in **Fig.SI9.3B**. The position of the maxima of the fringes, up to $k \simeq 40$ is found by fitting the radial profile to a multi-peak Gaussian function and the positions $x_k$ for which $\Delta\Phi(z_k, x_k) = 2\pi k$ (k=1,2…) can be compared to the one corresponding to the theoretical design.

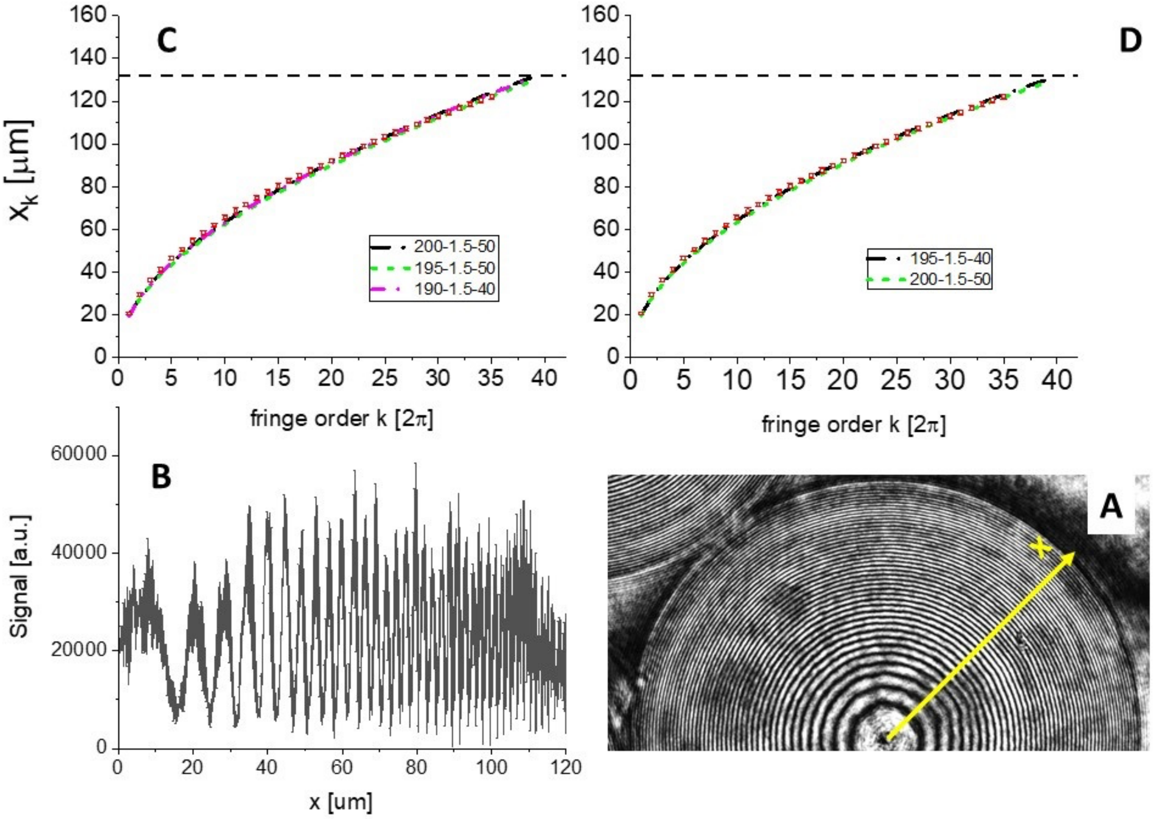


**Figure SI9.2: Panel A**: on axis digital holography of the microlenses displaying the phase shifts across the wavefront at a distance $d \simeq 50\mu m$ from the entrance pupil. **Panel B**: profiles of the image in (A) along the radial, x, direction averaged over 4 different polar angles. **Panel C**: comparison of the position of the fringes as a function of the order k of the fringes ($x_k$, open squares) compared with the theoretical prediction for a perfect spherical shape of the curved surface of the microlens (index of refraction n=1.5) for sphere radius $R = 200, 195\ and\ 190\ \mu m$. **Panel D**: comparison of the position of the fringes as a function of the order k of the fringes ($x_k$, open squares) compared with the theoretical prediction for a parabolic approximation of the curved sphere of the microlens (index of refraction n=1.5) for sphere radius $R = 200\ and\ 195\ \mu m$.

We have sampled the profile of the microlens on a plane at $50\ \mu m$ from the front vertex of the microlens, covering the whole size of the microlens (radius = $132\ \mu m$). The iso-OPD lines can be found by following the scheme in **Fig. SI9.3**. The total intensity on the CMOS camera is given by:

$$I_{tot}(r) = |E_{ref}(r,d_1) + E_{\mu lens}(r,d_2)|^2 = |E_{ref}(r,d_1)|^2 + |E_{\mu lens}(r,d_2)|^2 + 2Re\{E_{ref}E_{\mu lens}{}^*\}$$

(**SI9.1**)

The first two terms on the right-most hand side are the intensity of the reference (collimated) laser beam and the incoherent image of the microlens in transmission. The third term corresponds to the interference of the two beams. The two beams have propagation distance $d_1$ and $d_2$ along the top and bottom path of the Mach-Zender interferometer (**Fig.SI9.1**). If we write $E_{\mu lens}(r,d_2)$ $=E_{ref}(r,d_1)\ e^{k(d_1-d_2)}e^{i\Phi(r)}$ , we can rewrite the third term in **Eq. SI9.1** as:

$$2Re\{E_{ref}E_{\mu lens}{}^*\} = 2\ |E_{ref}(r,d_1)|^2 cos(k[d_1-d_2] + \Phi(r)) \qquad \textbf{(SI9.2)}$$

Although we cannot measure from **Eq. SI9.2** the full shape of the microlens, we can retrieve the iso-OPD lines that corresponds to the cases $\Phi(r_k) = 2\pi k$. The term $k[d_1-d_2]$ in **Eq. SI9.2** dictates only the intensity at the vertex of the microlens.

The condition $\Phi(r_k) = 2\pi k$ depends on the thickness of the microlens and index of refraction (n). It can be cast in terms of the shape of the microlens by referring to the sketch in **Fig. SI9.3**, as detailed hereafter. The phase shift of the ray impinging on the curved surface at a distance $\xi$ from the axis undergoes a phase shift up to the plane $\Pi_{obs}$ given by:

$$\Phi(x,z(x)) = k\ z(x) + kn\ C\bar{B} = k\ z(x) + kn\ [T-z(x)]\ /\ cos(\alpha-\beta) \qquad \textbf{(SI9.3)}$$

Since $tan(\alpha) = x_k\ /[R-z(x_k)]$ and $sin(\alpha) = n\ sin(\beta)$, we can write the phase of the beam passing through the lens up and emerging from the plane $\Pi_{obs}$ as a function of x and z(x). Moreover, by subtracting the phase $\Phi(x,z(x))$ to the constant reference value $\Phi_0 = nkT$, we can write that the phase difference is:

$$\Delta\Phi(x,z(x)) = knT\ [1-1/\ cos(\alpha-\beta)\ ] - k\ z(x)\ [1-n\ /\ cos(\alpha-\beta)] \qquad \textbf{(SI9.4)}$$

The condition to find the iso-OPD lines becomes therefore ($\Delta\Phi(x,z(x)) = 2\pi k$ ):

$$k\lambda = nT[1-1/cos(\alpha-\beta)] - z\ (x_k)[1-n/cos(\alpha-\beta)] \qquad \textbf{(SI9.5)}$$

k = 1,2,.....

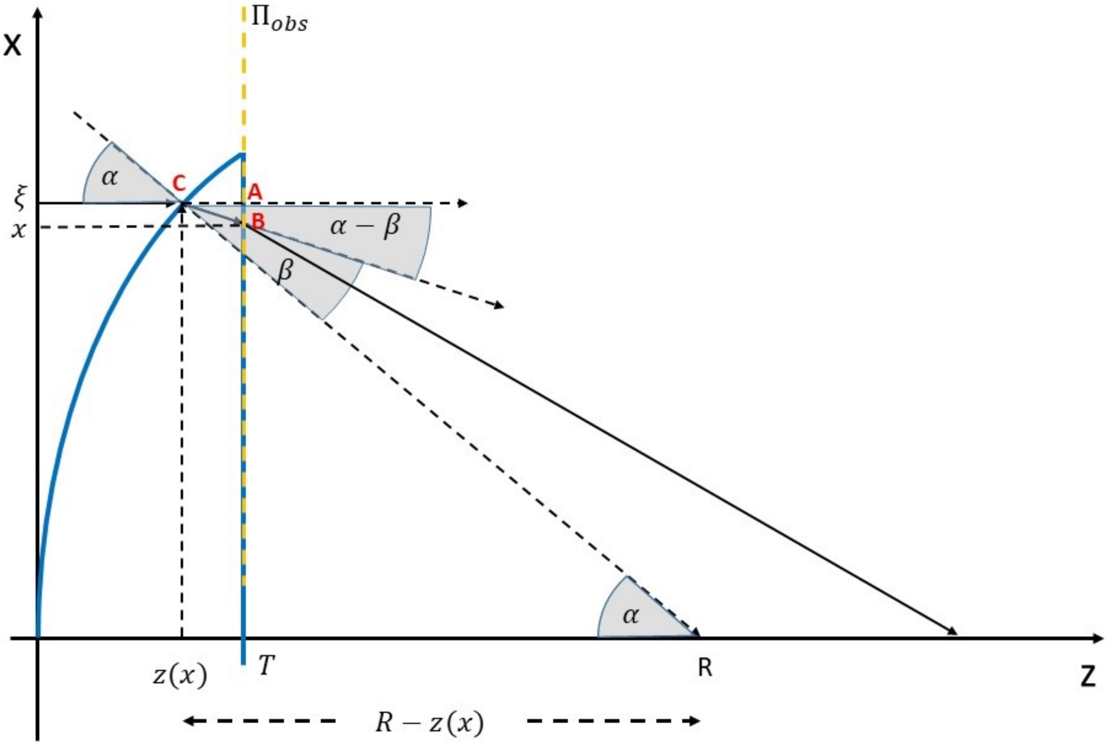


**Figure SI9.3**. Sketch of th refraction through the microlens and the computation of the iso-OPD lines ($\Phi(r_k) = 2\pi k$). Only the x coordinate is shown here, even though the problem has polar symmetry.

The position $x$ on the observation plane is related to the impinging position, $\xi$, by the relation:

$$x_k \to \xi_k - [T - z(x_k)tan(\beta)] \quad \textbf{(SI9.6)}$$

The pattern of the fringes deriving from the term $cos(k[d_1 - d_2] + \Phi(r))$ in **Eq. SI9.2** is clearly visible in the acquired images (**Fig.SI9.3A**). We are now in a position to predict the positions $x_k$ of the iso-OPD lines for a generic surface shape. For a perfect spherical shape (sag function, $z = R - \sqrt{R^2 - x^2}$) of the curved surface of the microlens and for the approximation $z \simeq x^2/(2R)$, which is the one adopted by Zemax. As one can judge from **Fig.SI9.2C-D**, we find in general an excellent agreement between theoretical prediction and experimental values for values of the radius $R \simeq 195\ \mu m$ and for $n = 1.5$, in agreement with the design of our microlenses. We can single out a slightly larger discrepancy between the experimental values and the theoretical predictions for the case of the perfect spherical shape (**Fig. SI9.2D**). The best agreement appears to occur for the case $z = \frac{x^2}{2R}\ R = 195\ \mu m$, assuming n=1.5.

#### SI10. UV irradiation Optimization during Fabrication Process.

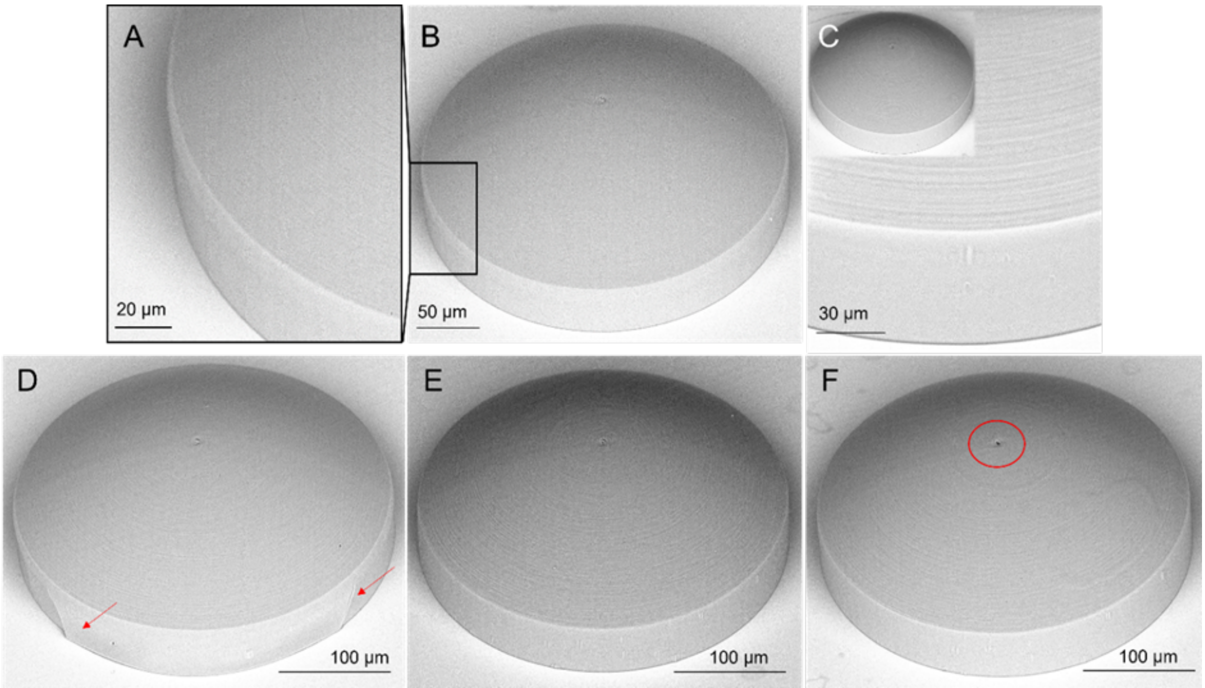


**Figure SI10:** Highlights on the laser powers and Uv exposure optimization procedures: B: Microlens fabricated using a laser power lower than the suitable one (13-15 mW with respect to the resist drop height); A: Highlight of the weak pedestal due to the not enough power used to polymerize it; C: Microlens directly UV exposed using source power of 200 mW for 120 sec. High power caused resin degradation, for instance the external surface became stiff turning yellow; D, E, F: UV exposure passing through the glass substrate; low UV power are not enough to crosslink the microlens inner core causing cracks in the lens pedestal (D, red arrows). Higher UV powers promote the distention of the outer surface improving the microlens smoothness. Too high power like 440 mW caused the formation of a central hole due to the excessive expansion of the inner resist (F, red ring). While, 320 mW is the best and suitable value to achieve the complete crosslink of the lens core avoiding any dome damage (E).
